# Structures of H2A.Z-associated human chromatin remodelers SRCAP and TIP60 reveal divergent mechanisms of chromatin engagement

**DOI:** 10.1101/2024.07.30.605802

**Authors:** Giho Park, Avinash B. Patel, Carl Wu, Robert K. Louder

## Abstract

H2A.Z is a conserved histone variant that is localized to specific genomic regions where it plays important roles in transcription, DNA repair, and replication. Central to the biochemistry of human H2A.Z are the SRCAP and TIP60 chromatin remodelers, homologs of yeast SWR1 which catalyzes ATP-dependent H2A.Z exchange. Here, we use cryo-electron microscopy to resolve six structural states of the native SRCAP complex, uncovering conformational intermediates interpreted as a stepwise path to full nucleosome engagement. We also resolve the structure of the native TIP60 complex which consists of a structured core from which flexibly tethered chromatin binding domains emerge. Despite the shared subunit composition, the core of TIP60 displays divergent architectures from SRCAP that structurally disfavor nucleosome engagement, suggesting a distinct biochemical function.

## Introduction

Chromatin plays a critical role in orchestrating essentially all genomic processes, from transcription to DNA repair and replication^1,2^. The basic repeating unit of chromatin, the nucleosome, consists of 146 base pairs (bp) of DNA wrapped around eight histone proteins, two copies of each of the canonical histones H2A, H2B, H3, H4^3,4^. While canonical histones are incorporated into chromatin in a replication-coupled manner, variant histones are expressed independent of the cell cycle and are dynamically incorporated through specific chaperone proteins or ATP-dependent chromatin remodeling complexes^5,6^. H2A.Z is a highly conserved histone variant involved in diverse chromatin functions, with dysregulation leading to neurological disease, cancer, and developmental defects^5–8^. Central to the specific deposition and post-translational modification of H2A.Z are the SRCAP and TIP60 complexes, respectively^9–12^. Both are homologs of yeast SWR1 which has been shown to exchange canonical H2A for H2A.Z in an ATP-dependent manner^13^. The TIP60 complex additionally contains an acetyltransferase module shown to acetylate H2A.Z^11,12^. The SRCAP and TIP60 complexes are recruited genome-wide to nucleosome free regions (NFRs) of promoters and enhancers^10,14–16^. Accordingly, H2A.Z deposition and modification occur in nucleosomes adjacent to NFRs, with the promoter proximal ‘+1 nucleosome’ downstream of transcriptional start sites (TSSs) being the best-characterized target^14,17–19^. The molecular mechanism of NFR targeting and subsequent nucleosome engagement by the two complexes remain elusive. Here, we elucidate structures of the SRCAP complex and demonstrate how the complex undergoes stepwise transitions for full nucleosome engagement. We also solve the structure of the TIP60 complex and present a comparative analysis with SRCAP, uncovering the structural and functional divergence of these two H2A.Z-associated chromatin remodelers.

## Results

### Structures of SRCAP suggest stepwise pathway from linker DNA to nucleosome engagement

To investigate the endogenous SRCAP complex, we purified to homogeneity the native 12-subunit, 1.0 MDa assembly from a gene-edited K562 cell line (see Methods) (Figure S1A). Notably, the native SRCAP (and TIP60, see below) complexes co-purified with the endogenous H2A.Z-H2B dimer (Figures S1A and S8A), as observed previously^20,21^. Purified SRCAP was incubated with nucleosomes containing long linker DNA (106N32) in the presence of ATPγS and analyzed by cryo-electron microscopy (cryo-EM). Importantly, we eschewed further purification of the SRCAP-nucleosome complex, subjecting the heterogeneous sample to cryo-EM analysis. This enabled us to discern multiple distinct states of SRCAP within the same sample, which we interpret as a stepwise pathway from linker DNA-binding to nucleosome engagement (Figures 1A and S1-3). The predominant nucleosome engagement state resembles previous structures of SWR1-nucleosome complex with the ATPase stably engaged on SHL2 (SHL positions defined in Figures 4A and 4B) where SWR1 is catalytically active for histone H2A.Z exchange (Figures 1A, S4C, and S4D)^22,23^.

**Figure 1:**
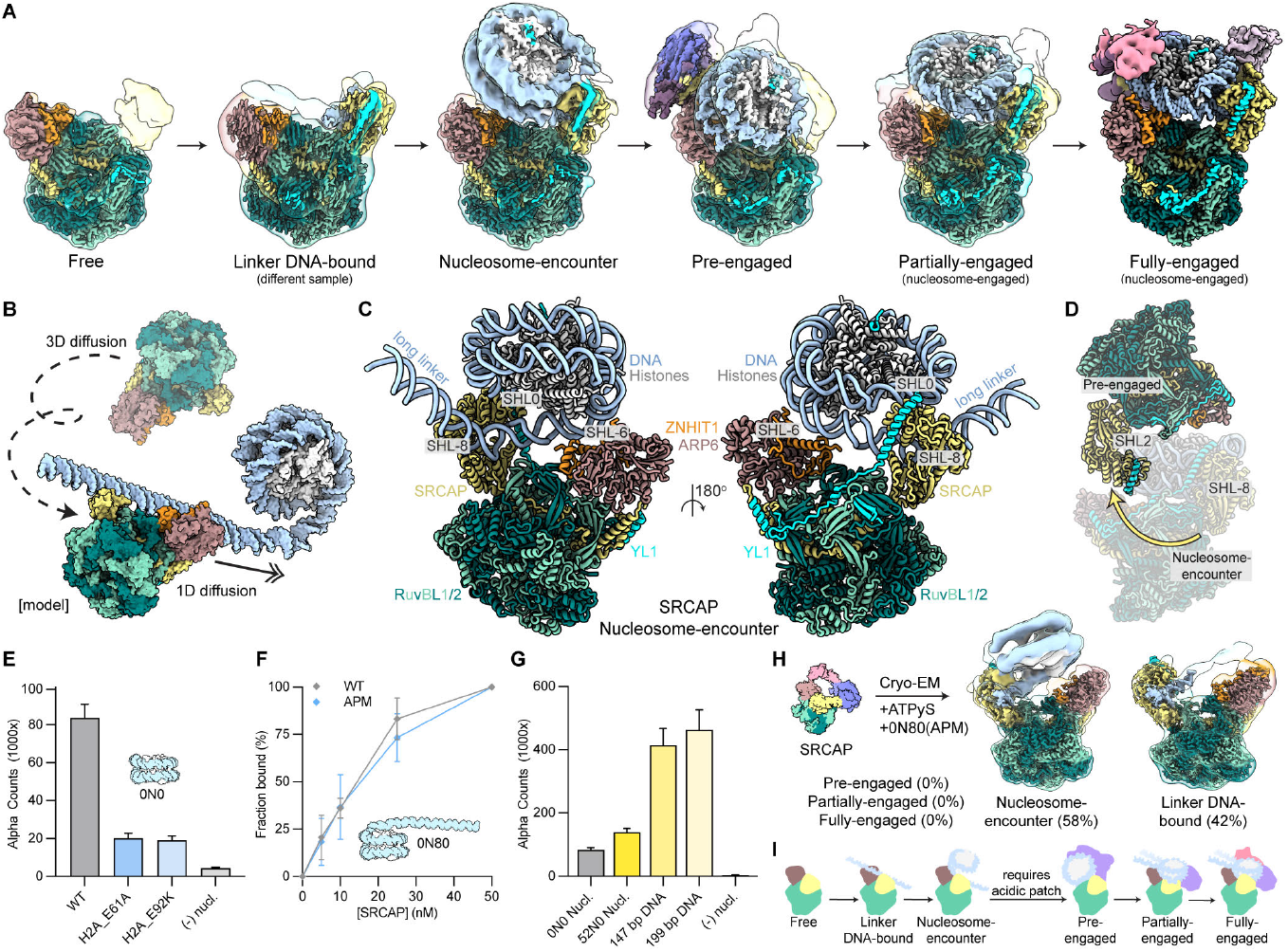
Structures of SRCAP suggest stepwise pathway from linker DNA to nucleosome engagement. **(A)** Cryo-EM reconstructions of the free, linker DNA-bound, nucleosome-encounter, pre-engaged, partially-engaged, and fully-engaged states of SRCAP. Low-pass filtered (transparent) and high-resolution composite (solid) maps are shown. **(B)** Conceptual model of SRCAP undergoing 3D diffusion to linker DNA after which it may undergo 1D diffusion to encounter a nucleosome. **(C)** Two views of the nucleosome-encounter state of SRCAP. The cryo-EM structure is shown as cartoon representation and major SHL contact sites are labeled. **(D)** The conformational flip in the transition from nucleosome-encounter state (transparent) to pre-engaged state. **(E)** AlphaScreen interaction assay results of SRCAP binding to WT, H2A_E61A, and H2A_E92K nucleosome core particles. Mean and standard deviation from three experiments are shown. **(F)** EMSA results of SRCAP binding to WT, APM (H2A_E61A/E64A/D90A/D92A) nucleosomes (0N80). Mean and standard deviation from three experiments are shown. **(G)** AlphaScreen interaction assay results of SRCAP binding to WT NCP (0N0), WT nucleosome (52N0), 147 bp naked DNA, and 199 bp naked DNA. Mean and standard deviation from three experiments are shown. **(H)** Results of the SRCAP-ATPγS-APM nucleosome cryo-EM dataset. Low-pass filtered (transparent) and high-resolution composite (solid) maps of SRCAP nucleosome-encounter and Linker-DNA bound are shown. **(I)** Cartoon depiction of the pathway from free to fully-engaged. The acidic patch is required to progress to states beyond nucleosome-encounter.

In the free state, most of the SRCAP complex is conformationally flexible and only the core module consisting of RUVBL1, RUVBL2, SRCAP, YL1, ZNHIT1, and ARP6 are visible (Figures 1A and S4E). This core with the exception of the SRCAP ATPase is structurally rigid and remains virtually unchanged in subsequent states. Notably, in addition to the monomeric free SRCAP complex, we observed dimeric SRCAP complexes in our sample (Figure S4F).

In the ‘nucleosome-encounter’ state, SRCAP complex is bound to the linker DNA adjacent to the nucleosome (Figure 1C). This state is strikingly distinct from previous structures of chromatin remodelers bound to nucleosomes and resembles the SRCAP complex having approached a nucleosome through 1D diffusion along the linker DNA (Figures 1B, 1C, and S4G)^24–29^. The complex is oriented ∼180° relative to the catalytically pertinent SHL2 position with the SRCAP ATPase engaged on linker DNA at SHL-8 while making secondary contacts at the nucleosome dyad (SHL0) (Figures 1C and 1D). Additionally, two positively charged regions (R^17^, R^26^, R^30^ and K^66^KKKKTR^72^) of the ZNHIT1 subunit dynamically interact with the nucleosome near SHL-6 (Figure S4H). Despite the flipped conformation of the nucleosome-encounter state, the YL1 subunit’s dual arginine anchors (R^130^, R^134^) bind to the H2A-H2B acidic patch as observed in subsequent states of the nucleosome engagement pathway (Figures 1C, S4I, and S4J). To further investigate the role of the YL1 acidic patch interaction in nucleosome engagement, we used the AlphaScreen assay to evaluate the binding of SRCAP complex to wild-type (WT) and acidic patch mutant (APM) nucleosome core particles (NCPs)^30^. Notably, single amino acid substitutions in the acidic patch (H2A_E61A or H2A_E92K) reduced the binding signal over four-fold, highlighting its critical role in SRCAP binding (Figure 1E). However, when we evaluated SRCAP complex binding to WT and APM (H2A_E61A/E64A/D90A/E92A) nucleosomes containing extended linker DNA (0N80), we observe no substantial differences in binding affinity, indicating that linker DNA binding by the SRCAP complex dominates over acidic patch interactions on the core histone surface (Figure 1F). We also observe a more than five-fold higher affinity for free DNAs (147 bp and 199 bp) over linker-containing nucleosomes (52N0) and NCPs, indicating the strong preference of the complex for free DNA in the range of NFR length (Figure 1G).

The biochemical data indicate that SRCAP complex interaction with the acidic patch is required for stable engagement with the nucleosome core. To explore this hypothesis structurally, we acquired a cryo-EM dataset with SRCAP bound to APM nucleosomes (0N80) in the presence of ATPγS (Figures 1H and S2A). We could identify linker DNA-bound and nucleosome-encounter states but did not observe any additional nucleosome bound states of the SRCAP complex (Figures 1H and S2A). In the linker DNA-bound state, SRCAP complex interacts with ∼40 bp of linker DNA through the SRCAP ATPase and the ZNHIT1 subunit (Figure S5A). The yeast SWR1 complex was shown to undergo 1D sliding on naked DNA and the structural similarity between the SWR1 and SRCAP DNA-bound complexes suggest a conserved mechanism of 1D diffusion on linker DNA leading to nucleosome engagement (Figures 1H, S5A, and S5B)^23,31^.

### SRCAP transition from pre- to nucleosome-engaged disrupts histone-DNA contacts

The Snf2-type ATPase of SRCAP is split into N-terminal and C-terminal lobes, between which a DNA-binding cleft is formed. We identified a distinct state of nucleosome-bound SRCAP in which the ATPase is contacting the catalytically pertinent SHL2 position, but without the DNA fully bound within the cleft (Figures 2A and 2C). In this state, the ATPase is splayed wide open and only the N-lobe interacts with SHL2 and -6 (Figures 2A and 2C). Additionally, a positively charged loop of ZNHIT1 (K^66^KKKKTRGDHFKLRFR^81^) contacts the H2A-H2B dimer near the acidic patch (Figure S5C). We term this state ‘pre-engaged’, as the ATPase has targeted SHL2 but is not yet engaged. Notably, in this state, the nucleosome is spatially distant from the SRCAP core module and the nucleosomal DNA remains fully wrapped (Figure 2A).

**Figure 2:**
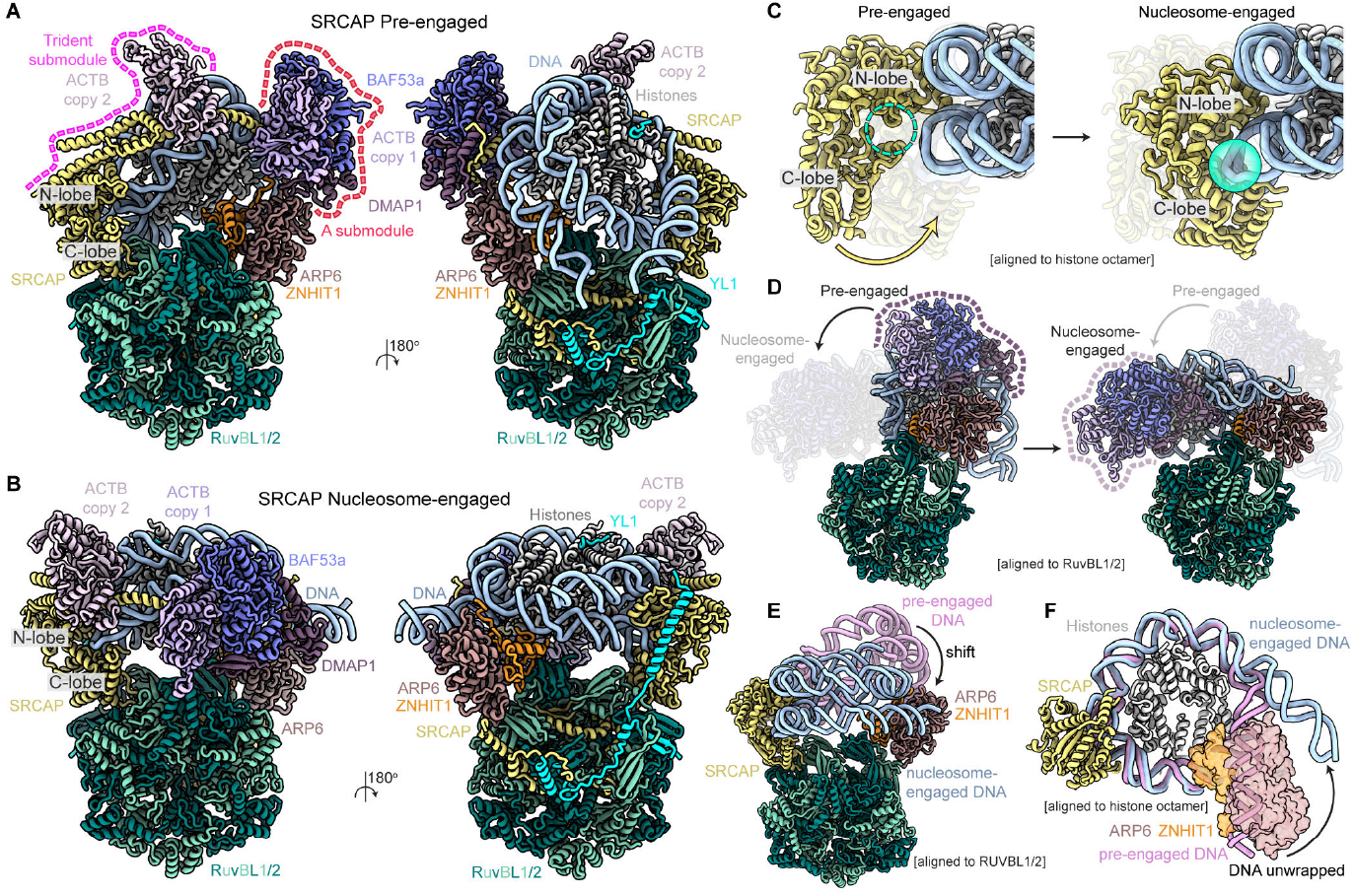
SRCAP transition from pre- to nucleosome-engaged disrupts histone-DNA contacts. **(A)** Two views of pre-engaged SRCAP-nucleosome complex shown as cartoon representation. The ‘Trident’ and “A’ submodules are indicated with pink and red dotted lines, respectively. **(B)** Two views (same views as panel A) of nucleosome-engaged SRCAP-nucleosome complex shown as cartoon representation. **(C)** Conformational shift of the SRCAP ATPase in the transition from pre-engaged to nucleosome-engaged. The N- and C-lobes of the ATPase are labeled. The empty and DNA bound ATPase cleft are labeled with a dashed and solid green sphere, respectively. **(D)** Conformational shift of the HSA module in the transition from pre-engaged to nucleosome-engaged. Some subunits are not shown for clarity. The two states were aligned to the RUVBL1/2 core. **(E)** Conformational shift of the pre-engaged nucleosome (pink) to the nucleosome-engaged nucleosome (blue). The two states were aligned to the RUVBL1/2 core. **(F)** During the transition from pre-engaged (pink) to nucleosome-engaged (blue), the ARP6-ZNHIT1 subunits wedge between the histone core and unwrap ∼20 bp of DNA. Some subunits are not shown for clarity and the two states were aligned to the histone octamer.

Beyond the SRCAP core module, the N-terminal 255 amino acids of SRCAP subunit includes the HSA (Helicase/SANT-Associated) domain which serves as a structural scaffold for the assembly of a 250 kDa module composed of GAS41, DMAP1, BAF53a, and two copies of Actin (Figures 2A, 4A, S7C, and S7D). Strikingly, in the pre-engaged state and all subsequent states, the SRCAP HSA extended alpha-helix arcs ∼80 degrees to envelop the nucleosome (Figures 2A, 4A, and S7D). This ‘HSA’ module is divided into three structural submodules, two of which are observed in the pre-engaged state (Figures 2A and S7C). The first ‘trident’-shaped submodule includes the C-terminal end of SRCAP HSA helix which extends outward from the ATPase and interacts with Actin (copy 2) and another conserved C-terminal segment of the SRCAP subunit (Figures 2A, 4A, 4B, and S7C). The second ‘A’ submodule consists of the extended HSA of SRCAP, Actin (copy 1), BAF53a, and DMAP1 (Figures 2A, 4A, and S7C). In the pre-engaged state, the N-terminus of the DMAP1 subunit is bound by ARP6 through a hydrophobic pocket that accommodates L86 of DMAP1 (Figures S5D and S5E). This DMAP1-ARP6 interaction is disrupted upon nucleosome-engagement.

Given that the two lobes of the SRCAP ATPase in the pre-engaged state exhibit a more ‘open’ configuration compared to the nucleosome-engaged state, we speculated that nucleotide binding facilitates engagement. A cryo-EM dataset of SRCAP-nucleosome complex assembled in the absence of nucleotide contained only nucleosome-encounter and pre-engaged states (Figure S2C), confirming that the transition from the pre-engaged state to the nucleosome-engaged state is contingent on the presence of nucleotide.

In the subsequent nucleosome-engaged state, the SRCAP ATPase cleft is stably bound to SHL2 and the HSA module has undergone a significant conformational change (Figures 2B, 2C, and 2D). Additionally, the nucleosome is positioned closer to the RUVBL1/2 core and ∼20 bp of DNA are unwrapped near SHL6 to SHL7 (Figures 2E and 2F). Presumably, engagement of the ATPase cleft of SHL2 DNA induces conformational changes of the connected HSA module. Along with ARP6 release of DMAP1, the HSA module and nucleosome are able to spatially shift closer to the core module to attain the nucleosome-engaged position. During this transition the nucleosome is physically wedged by the ZNHIT1-ARP6 subunits, detaching ∼20 bp of DNA from the histone octamer (Figures 2E, 2F, and S5F). Accordingly, we propose that SRCAP transitions from the pre-engaged state to the nucleosome-engaged state as a mechanism to disrupt histone-DNA contacts independent of ATP hydrolysis.

### Structural basis for nucleosome engagement by SRCAP core module

The high-resolution (2.6-3.2 Å) map of the nucleosome-engaged SRCAP core module enabled us to accurately model the structure at the atomic level (Figure 3A). Overall, the human complex is organized similarly as previously reported yeast SWR1-nucleosome structures^22,23^, featuring the two key nucleosome binding modules, SRCAP/YL1 and ZNHIT1/ARP6, positioned on opposite sides of the RUVBL1/2 hetero-hexamer hub, into which the SRCAP insert domain is incorporated (Figure 3A). Further classification of the cryo-EM data reveals the partially- and fully-engaged states, with the latter state forming more extensive interactions between the ZNHIT1/ARP6 module and the nucleosome (discussed below).

**Figure 3:**
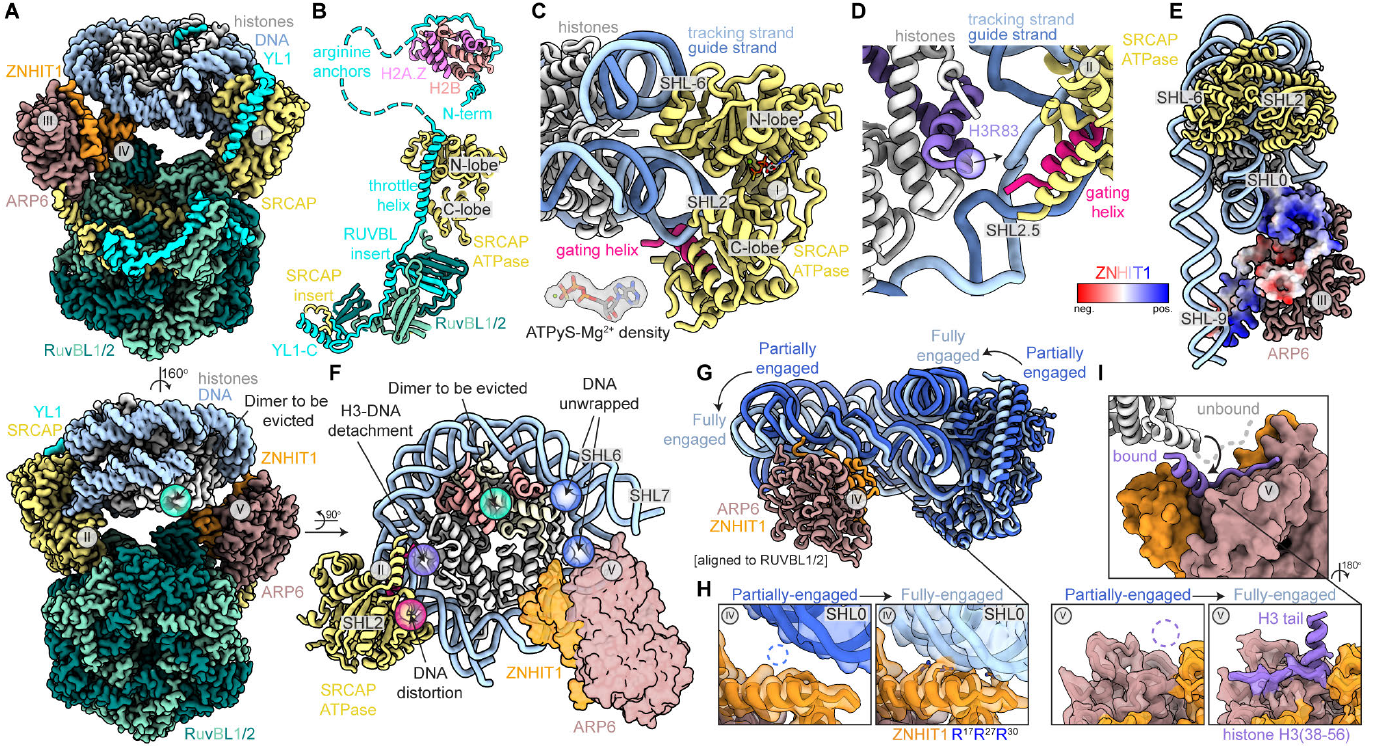
Structural basis for nucleosome engagement by SRCAP core module. **(A)** Two views of the cryo-EM composite map of the core fully-engaged SRCAP-nucleosome complex. Roman numerals are used to label different regions of the complex discussed in subsequent panels. The H2A-H2B dimer destined for eviction is labeled with a green sphere. **(B)** Isolated view of YL1 subunit organization. All elements are modeled from cryo-EM data except for the YL1-H2A.Z-H2B subcomplex crystal structure (PDB: 5FUG). Disordered loops are indicated with dashed lines and some subunits are not shown for clarity. **(C)** Close-up of SRCAP ATPase (region I) that binds primarily to SHL2 and forms additional contacts at SHL-6. The N- and C-lobe of the ATPase are labeled and the gating helix is colored pink. The tracking and guide strand of the nucleosome DNA are colored light and dark blue, respectively. The cryo-EM density of ATPγS-Mg^2+^ in the N-lobe nucleotide pocket is shown. Atoms are colored by elements (red: oxygen, blue: nitrogen, orange: phosphate, green: magnesium). **(D)** Close-up of the gating helix (region II) that inserted into the minor groove near SHL2.5 and distorts the DNA, disrupting histone H3 contacts (purple sphere). **(E)** Side-view showing multivalent interactions of ZNHIT1 (region III) with the entry DNA near SHL-9 and dyad DNA at SHL0. ZNHIT1 is colored according to Coulombic electrostatic potential (red: -10 kcal/(mol·e), blue: +10 kcal/(mol·e)). **(F)** Isolated view of the nucleosome, ATPase, and ZNHIT1/ARP6 module. DNA distortion in the ATPase cleft, DNA detachment from H3, H2A-H2B dimer to be evicted, and the unwrapped DNA are labeled with pink, purple, green, and blue spheres, respectively. **(G)** Isolated view of the nucleosome, ATPase, and ZNHIT1/ARP6 module showing the conformational change from partially-engaged (dark blue) to fully-engaged (light blue). The two states were aligned relative to the RUVBL1/2 core. **(H)** Zoom in views of ZNHIT1 (region IV) showing dyad (SHL0) DNA binding in the fully-engaged state. The cryo-EM map is shown as transparent and the model is shown as a cartoon representation. Atoms are colored by elements (blue: nitrogen). **(I)** Histone H3 tail binding by ARP6 (region V) is shown with unbound H3 colored white and bound H3 colored purple (top). Rotated and zoomed in views (bottom) show H3 tail binding in the fully-engaged state. The cryo-EM map is shown as transparent and the model is shown as a cartoon representation.

**Figure 4:**
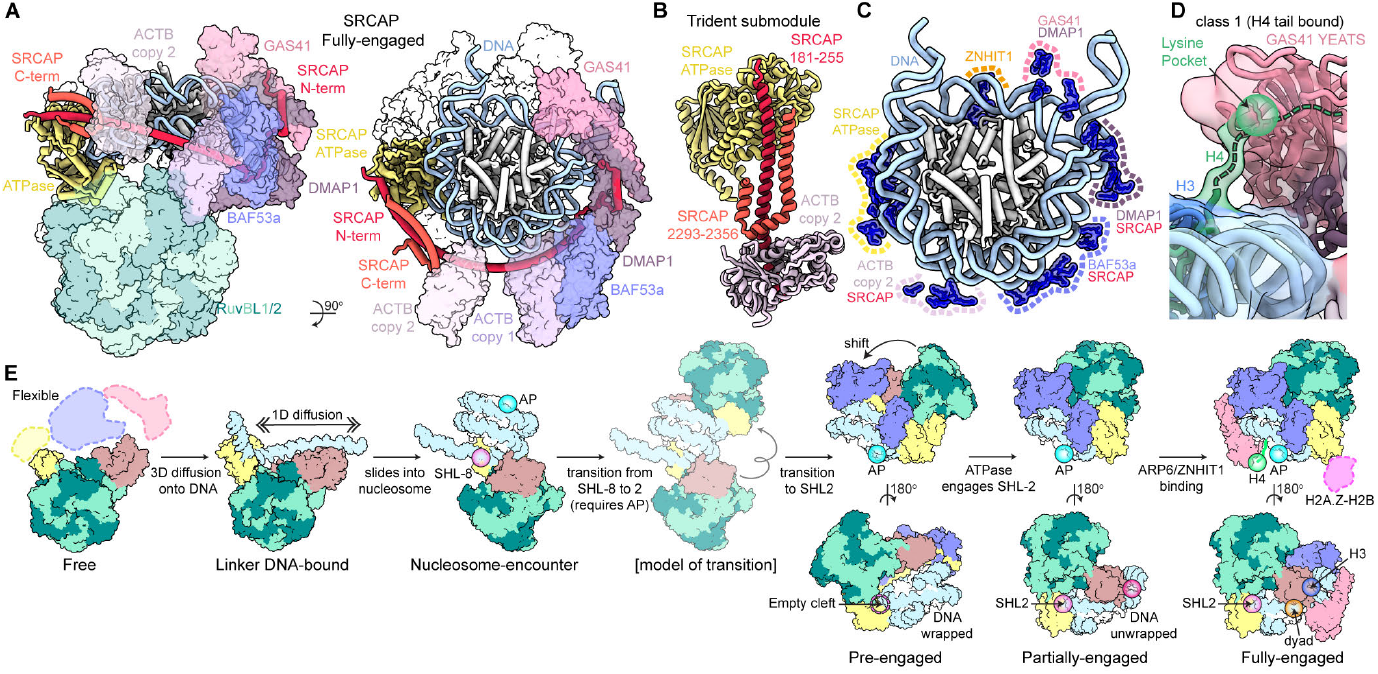
SRCAP envelops nucleosome on full engagement. **(A)** Complete cryo-EM structure of fully-engaged SRCAP-nucleosome complex in two views. Regions of SRCAP and nucleosome are shown as ribbon representation and other subunits as transparent surfaces. SRCAP ATPase, N-terminal HSA helix, and C-terminal region are colored yellow, red, and orange, respectively. **(B)** Isolated view of the ‘Trident’ submodule with SRCAP ATPase, N-terminal HSA helix, and C-terminal region colored yellow, red, and orange, respectively. **(C)** Lysine and arginine atoms of SRCAP that likely contact the nucleosome DNA are shown. The submodules and the corresponding subunits are indicated with dashed lines. **(D)** Cryo-EM map of the H4 tail bound class is shown with the lysine pocket of GAS41 YEATS domain highlighted green. The H4 tail is shown as a dashed green line. **(E)** Cartoon depiction of stepwise nucleosome engagement by SRCAP.

Our model of the SRCAP core module reveals the critical role of the YL1 subunit in organizing the complex (Figure 3B). The N-terminal region of YL1 contains the H2A.Z-H2B chaperone domain for which the crystal structure has been previously determined^32,33^. We could putatively dock the YL1-H2A.Z-H2B structure into low resolution cryo-EM density adjacent to the SRCAP ATPase (Figures S5G and S5H). Notably, the catalytic SRCAP subunit contains an additional H2A.Z-H2B chaperone domain adjacent to the N-lobe of the ATPase which may function cooperatively with the YL1 chaperone domain (Figures S5G, S5H, and S5I)^34^. Our docking reveals the close proximity of H2A.Z-H2B with the SRCAP ATPase and SRCAP HSA helix, suggesting a coupling between ATP hydrolysis and the release of the H2A.Z-H2B dimer for histone exchange (Figures S5H and S5I)^35^.

As observed in previous states, YL1 contains a dual arginine anchor motif that interacts with the H2A-H2B acidic patch on the nucleosome face opposite to the H2A-H2B dimer that is targeted for exchange (Figures S4I and S4J) This is followed by a linker that passes over SHL-6 and connects the arginine anchor to the YL1 throttle helix. The YL1 throttle bridges the two N- and C-lobes of the SRCAP ATPase and is followed by a beta-hairpin that inserts between the OB folds of RUVBL1 and -2, constraining the SRCAP ATPase relative to the core. The carboxyl-terminus of YL1 harbors the highly conserved YL1-C domain. In yeast, the YL1-C domain of Swc2 forms a subcomplex with the fungal-specific Swc3 subunit that was shown to be critical for SWR1 chromatin binding and promoter-specific H2A.Z deposition genome wide (Figures S6A and S6C)^23^. In the human complex, YL1-C instead folds with the insert domain of the SRCAP subunit (residues 987 to 998) (Figures 3B and S6B). While the core YL1-C fold is evolutionarily conserved, the human YL1-C:SRCAP submodule lacks the structured DNA binding domains observed in yeast Swc2:Swc3 submodule (Figures S6A-C). Rather, the SRCAP subunit contains a disordered 900 amino acid loop (residues 999 to 1881) that extends from the complex (Figure S6D), suggesting the evolution of a unique, intrinsically disordered region (IDR)-based mechanism for target search, as observed for the cBAF complex^36^. Notably, the C-terminus of SRCAP subunit contains another large disordered region (residues 2357 to 3230) that contains three AT hooks shown to be important for nuclear localization and nucleosome binding of SRCAP, with mutations in this region leading to Floating-Harbor Syndrome (Figure S6D)^37^.

The SRCAP ATPase primarily binds the nucleosomal DNA at SHL2 through its cleft between the N- and C-lobes (Figure 3c). In addition, the N-lobe is inserted between the two gyres of nucleosomal DNA, establishing secondary contacts at SHL-6 (Figure 3c). The binding of the ATPase distorts the nucleosomal DNA, inducing a large bulge of both the tracking and guide strands of DNA (Figure S6E). The DNA is further distorted by the gating helix which inserts into the minor groove of the bulge (Figure 3D). Relative to previously investigated remodelers, the SRCAP gating helix is inserted deeper into the widened minor groove, disrupting DNA contacts to H3 loop 1 (R^83^) at SHL2.5 and inducing a kink (Figures 3D and S6J)^38–42^. The observed DNA distortion and detachment from histones occurs on the side of the nucleosome where the DNA is unwrapped for H2A-H2B dimer eviction, indicating that the fully engaged SRCAP is poised for exchange (Figure 3F). Notably, despite the unambiguous presence of ATPγS in the nucleotide pocket, the SRCAP ATPase resembles the “open” or apo state of other chromatin remodelers (Figures 3C and S6G-J)^38–42^. These differences in nucleotide-conformational coupling may arise from the unique histone exchange activity of SRCAP, which does not involve translocation of the nucleosomal DNA.

Upon transition from the partially- to fully-engaged state, the ZNHIT1/ARP6 module forms extensive interactions with DNA and histone H3, bringing the nucleosome closer to core SRCAP (Figures 3E and 3G-I). The N-terminal alpha-helix of ZNHIT1 (R^17^, R^26^, R^30^) binds to nucleosomal DNA at the dyad (SHL0) (Figures 3G and 3H). Moreover, a positively charged loop (K^66^KKKKTR^72^) of ZNHIT1 dynamically interacts with the linker DNA near SHL-9 (Figures 3E and S6K). ARP6 binds the unmodified histone H3 N-terminal tail (residues 38-56), with H3 Y^41^ inserting into a hydrophobic pocket on the ARP6 surface (Figures 3I, S5D, and S7A). The interaction appears to be independent of histone post-translational modifications and likely functions in anchoring to the histone octamer and sequestering the positively charged tail to prevent re-wrapping of the nucleosomal DN^43^. The H3-ARP6 interaction is dynamic as we were unable to fully separate H3 bound and unbound populations within the particle subset corresponding to the fully-engaged state (Figure S7B). Collectively, these interactions highlight the function of the ZNHIT1/ARP6 module in disrupting histone-DNA contacts and firmly anchoring the nucleosome, facilitating its full engagement by the SRCAP complex.

### SRCAP envelops nucleosome on full engagement

In the fully-engaged state, we observe the full HSA module including the third submodule which forms a tetrameric coiled-coil structure with DMAP1, GAS41, and the N-terminus of the SRCAP subunit (Figures 4A, S7C, and S7D). Remarkably, the full SRCAP complex envelops the nucleosome and establishes extensive electrostatic contacts with the DNA through its various subunits (Figures 4C, S7D, and S7E). The SRCAP ATPase binds to SHL2 and -6. ZNHIT1 binds to the dyad at SHL0. The trident submodule contacts SHL3 and SHL-4. The A submodule contacts SHL4 to 5. SRCAP and DMAP1 contact SHL5 and 6. Finally, the GAS41 and DMAP1 contact SHL-2 to -1.

Intriguingly, within the nucleosome-bound complex, GAS41 is positioned near the histone H4 N-terminal region (Figures 4A, 4D, and S7G). Further classification of the cryo-EM data revealed a state wherein the YEATS domain of GAS41 binds to the unmodified H4 tail (Figure 4D). While we can trace the H4 N-terminal tail from the nucleosome core to the lysine pocket and speculate that the unmodified H4K16 is most likely bound (Figure S7F), we cannot rule out more N-terminal lysines such as H4K5/8/12. This finding contrasts with earlier studies concluding that GAS41 preferentially binds to the H3 N-terminal tail^44–47^. However, those studies used the YEATS domain or GAS41 subunit in isolation and all but one study probed histone peptides, rather than the nucleosome substrate^44–47^. While we cannot rule out the possibility of a transient H3 interaction in an unobserved state from the cryo-EM datasets, our findings underscore the importance of investigating histone PTM readers within the context of their full complexes and with nucleosomal substrates^48,49^.

Collectively, our structures define a stepwise pathway of nucleosome engagement by the SRCAP complex (Figure 4E). As the complex transitions through the linker DNA-bound, nucleosome-encounter, pre-engaged, partially-engaged, and fully-engaged states, it adopts unique structural conformations and interacts with the DNA and histone proteins in distinct configurations (Figure 4E). The structure of the fully-engaged state includes all 17 polypeptides of the SRCAP complex including the H2A.Z-H2B dimer and likely represents an enzymatically poised state just prior to histone exchange (Figure 4E).

### Architecture of TIP60 and model for chromatin binding

TIP60 is a 19-subunit 1.85-MDa complex that plays diverse roles in acetylation of both histone and non-histone proteins through the catalytic TIP60 subunit (also known as KAT5), thus affecting the overall structure of chromatin, regulating the binding of effector proteins, and modulating functions of non-histone proteins^50^. Furthermore, TIP60 has been proposed to catalyze histone H2A.Z exchange though the Snf2-type ATPase containing EP400 subunit^16,51–53^. Despite the widespread functional implications of TIP60, there is no structural information available for the native human TIP60 complex other than a crystal structure of the dimeric MBTD1-EPC1 subcomplex^54^. We purified the native TIP60 complex in a similar manner to the native SRCAP complex (see Methods) and analyzed the sample using electron microscopy (Figures S8A, S9A, S9B).

Initial 2-D negative stain EM analysis of the endogenous TIP60 sample showed the complex to comprise two main lobes, with one lobe corresponding to the distinctive DNA-PK-like structure of the TRRAP subunit^30,55^, and the second lobe corresponding to the hexameric RuvBL1/2 hub (Figures 5A and 5B). These two lobes, which will be referred to throughout as TRRAP and RUVBL, exhibit an extremely high degree of flexibility with respect to one another (Figures 5A and 5C).

**Figure 5:**
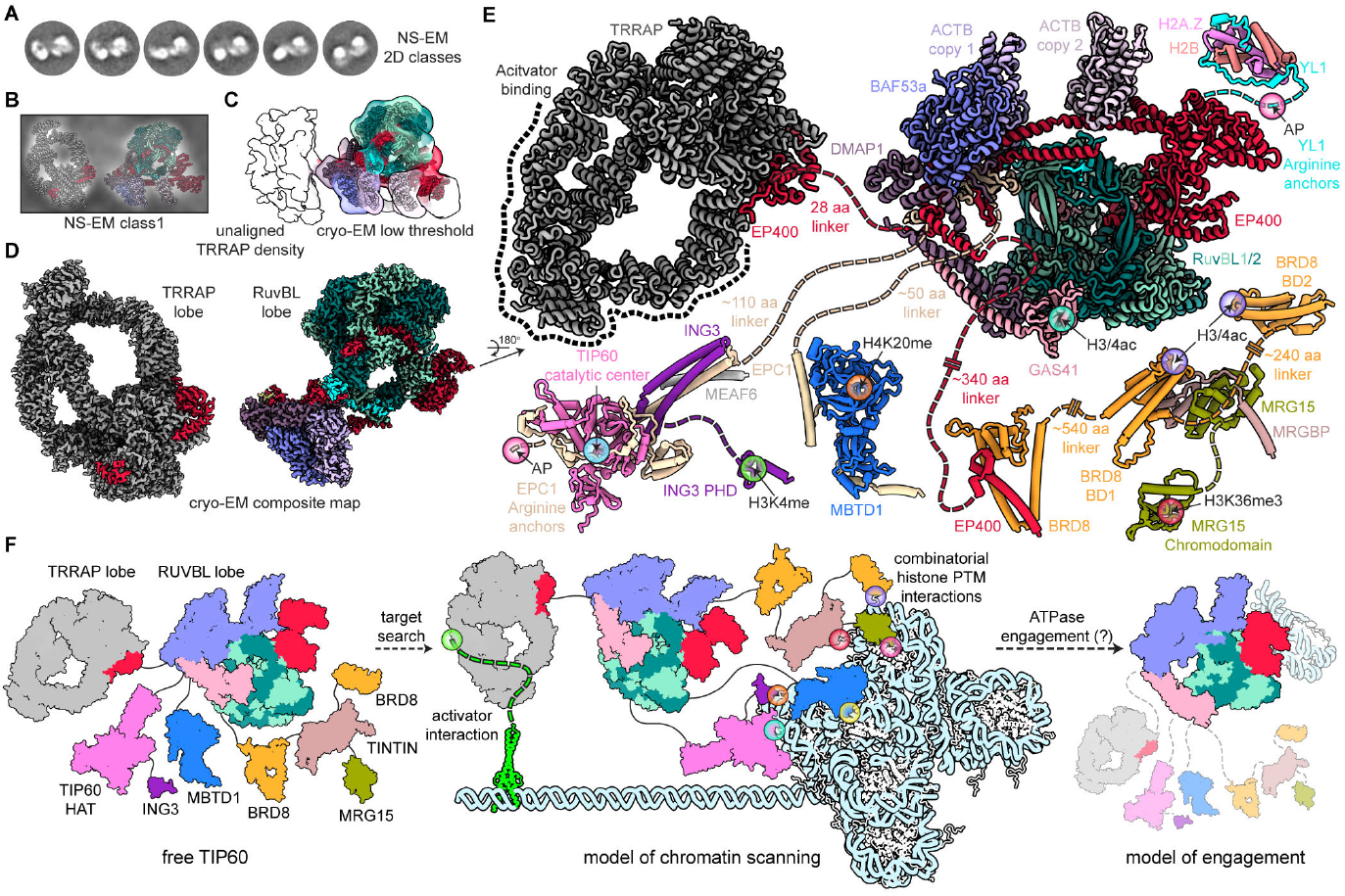
Architecture of TIP60 and model for chromatin binding. **(A)** Negative Stain (NS) 2D class averages of native human TIP60 complex. **(B)** Exemplar NS-EM 2D class showing the relative orientation between the TRRAP and RUVBL lobes. **(C)** Cryo-EM map of native human TIP60 complex at a low threshold showing the flexible unaligned TRRAP density. **(D)** Cryo-EM composite maps of the TRRAP and RUVBL lobes (orientated as in panel B). **(E)** Integrative model of TIP60 based on cryo-EM structures and AlphaFold2 multimer predictions (shown as tube helices). Dotted lines represent linker between structured domains. Post-translational modification (PTM) binding sites are highlighted with spheres. **(F)** Cartoon depiction of a model for nucleosome engagement by TIP60.

Cryo-EM analysis of native TIP60 resulted in reconstruction of the TRRAP and RUVBL lobes to 3.0-Å and 3.4-Å resolution respectively (Figures S8A, S9A, and S9B). To further probe the structural independence between the TRRAP and RUVBL lobes, we co-expressed the identified subunits from each lobe ectopically in human HEK293 cells (Figures S8B, S8C, S9C, and S9D). We purified the two subcomplexes and solved cryo-EM structures of the TRRAP and RUVBL lobes to 2.3 Å and 2.7 Å, respectively (Figures 5D, S8B, S8C, S9C, and S9D). The regions resolved by cryo-EM are essentially identical between the ectopically expressed and native TIP60 complexes, indicating the two lobes are organized independently of each other (Figures S8A-C and S9A-D). Analysis of the maps revealed that the TRRAP lobe contained the entirety of the 438 kDa TRRAP protein along with parts of EP400 (Figures 5D, 5E, and S10A-C). The RUVBL lobe is organized similarly to SRCAP (this study) and other remodelers of the INO80 family, with RUVBL1 and RUVBL2 forming a heterohexameric core that incorporates the EP400 insert domain (Figures 5D and 5E) ^27^. The EP400 Snf2-type ATPase emerges from the insert domain and is tethered to the core via C-lobe interactions with RUVBL1 and 2. Extending from the N-lobe, the EP400 extended HSA traverses towards the opposite side of the RUVBL1/2 hexamer where it scaffolds the HSA module composed of ACTB, BAF53a, DMAP1, EPC1, and GAS41. The C-terminal EP400 domain additionally interacts with the HSA module and connects the RUVBL and TRRAP lobes through a flexible 28 amino acid linker.

The HSA modules of TIP60 and SRCAP are structured similarly (Figures 4A and 5E). ACTB and BAF53a form a tight heterodimer on the EP400 HSA and DMAP1 anchors to the heterodimer through a conserved hydrophobic beta-hairpin and SANT domain interactions with BAF53a (Figure 6D). EP400 additionally forms a tetrameric coiled-coil structure with DMAP1 and GAS41, which we were able to putatively model using AlphaFold2 and lower resolution maps (Figures 5E and S9F). Unique to TIP60, the N-terminal region of DMAP1 interacts with the EP400/EPC1 beta cluster and the YL1-C fold (Figures 6D and 6E). Further, the C-terminus of EP400 stabilizes the HSA module through interactions with RUVBL1/2 and the YL1 beta-hairpin (Figure 6A). Overall, these interactions anchor the HSA module of TIP60 to the RUVBL core, an arrangement distinct from the flexibly attached HSA modules observed in the SRCAP (this paper) and INO80 complexes^27,42^.

**Figure 6:**
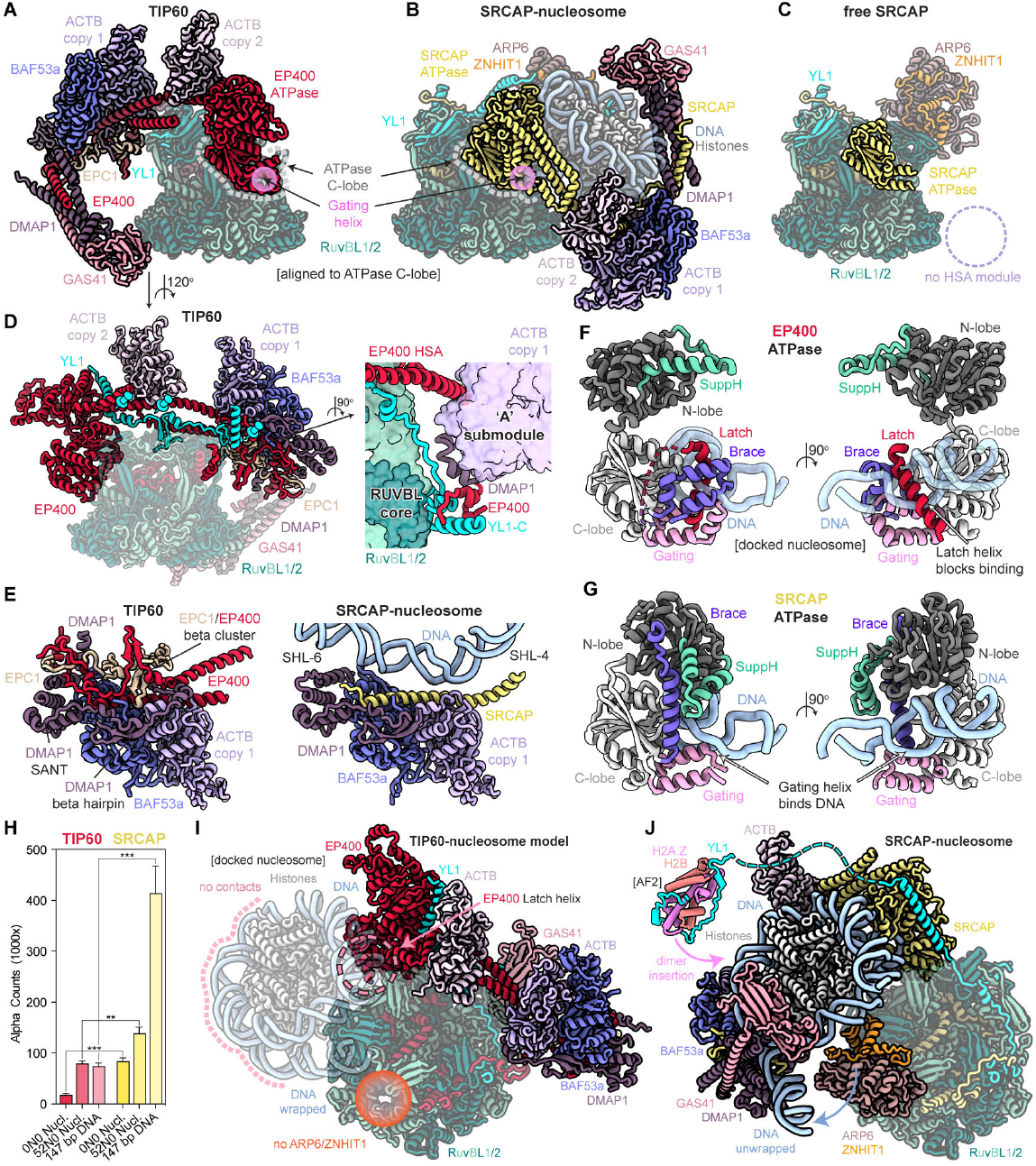
Structural divergence of SRCAP and TIP60. **(A)** Cryo-EM structure of TIP60 RUVBL lobe. The EP400 ATPase C-lobe is outlined with a gray dotted line and the gating helix highlighted with a green sphere. **(B)** Cryo-EM structure of SRCAP-nucleosome complex aligned to TIP60 ATPase C-lobe. The SRCAP ATPaseC is outlined with a gray dotted line and the gating helix highlighted with a green sphere. **(C)** Cryo-EM structure of SRCAP (free) aligned to TIP60 ATPase C-lobe. Dotted circles represent the absence of the HSA module (purple). **(D)** Rotated view of TIP60 RUVBL lobe structure (panel A). The elements of YL1 that anchor the HSA module to the core are highlighted with cyan pins. A zoomed-in view of the YL1-C interaction with EP400 and DMAP1 which anchor the ‘A’ submodule to the RUVBL core is shown (right). **(E)** Comparison of the TIP60 and SRCAP-nucleosome HSA module aligned to ACTB and BAF53a heterodimer. For SRCAP-nucleosome, the histones were removed for clarity. **(F)** Isolated views of the EP400 ATPase with docked nucleosomal DNA (as in panel H). The N-lobe, C-lobe, SuppH, Brace, Gating, Latch, and DNA are colored dark gray, light gray, green, purple, pink, red, blue, respectively. The unique Latch helix of EP400 blocks the Gating helix from binding DNA. **(G)** Isolated views of the SRCAP ATPase bound to nucleosomal DNA (fully-engaged state). The N-lobe, C-lobe, SuppH, Brace, Gating, and DNA are colored dark gray, light gray, green, purple, pink, blue, respectively. The SRCAP Gating helix inserts into the minor groove and detaches the DNA form the histone core. **(H)** AlphaScreen interaction assay results of TIP60 (red) and SRCAP (yellow, same as Fig. 1G) binding to WT NCP (0N0), WT nucleosome (52N0), and 147 bp naked DNA. Mean and standard deviation from three experiments are shown. Two-tailed, unpaired Student’s t-test was used for statistical analysis (***P = 0.0001 for 0N0 Nucl.; **P = 0.0013 for 52N0 Nucl.; ***P = 0.0004 for 147 bp DNA). **(I)** TIP60-nucleosome model based on docking the nucleosome from the fully-engaged SRCAP structure (panel I). The pink dotted lines indicate HSA module nucleosome contacts that are missing in TIP60. The pink dotted circle represents steric clash between the latch helix and the nucleosome. The orange sphere shows the absence of ARP6 and ZNHIT1 subunits which unwrap DNA in SRCAP. **(J)** Cryo-EM Structure of nucleosome fully-engaged SRCAP poised for histone exchange. The HSA module envelops the nucleosome through extensive electrostatic contacts. ARP6/ZNHIT1 unwraps approximately 20 base pairs of DNA. YL1 chaperones the H2A.Z-H2B dimer for exchange. The YL1-H2A.Z-H2B is modeled based on AlphaFold2 multimer prediction.

While our cryo-EM structures resolved the ∼600 kDa RUVBL lobe and ∼450 kDa TRRAP lobe, additional subunits of the TIP60 complex, largely corresponding to chromatin binding modules accounting for another ∼300 kDa, were not visible due to their flexible attachment to the main lobes. In all, disordered regions account for roughly 500 kDa, or 26% of the 1.85 MDa complex. To understand how the entire complex is organized, we utilized AlphaFold2 to generate predicted structures of the missing modules (Figures 5E, S9E, S9G, and S9H)^56–58^. The N-terminus of the EPC1 subunit associates with the TIP60 lysine acetyltransferase (KAT) along with ING3, and MEAF6 (Figure 5E). This KAT module is connected to the HSA module through a ∼130 amino acid unstructured linker, indicating a flexible attachment conducive to long-distance acetylation (Figure 5E). The C-terminus of EPC1 folds with the MBTD1 subunit (Figure 5E)^54^. The C-terminus of MRG15 associates with MRGBP through its MRG domain and MRGBP further interacts with bromodomain 1 of BRD8 to form the conserved TINTIN module (Figure 5E)^59^. The N-terminal region of EP400 tethers BRD8 on a long ∼340 amino acid linker (Figure 5E). Notably the various epigenetic modification binding modules are tethered on long linkers, enabling TIP60 to scan distant nucleosomes within the chromatin environment (Figure 5F). We speculate that the combinatorial actions of epigenetic modification binding modules, Snf2-type ATPase of EP400, and activator interactions with TRRAP^60^ provide a stable interaction for TIP60 recruitment to promoters and enhancer (Figure 5F).

### Structural divergence of SRCAP and TIP60

Nine subunits are shared between the SRCAP complex and RUVBL lobe of the TIP60 complex, with ZNHIT1 and ARP6 being unique to SRCAP, and EPC1 being unique to TIP60. The two complexes share a near identical structure of the RUVBL1/2 hexamer, EP400/SRCAP insert domain, and YL1 (Figures 6A-C and 6E). However, despite a near-identical subunit composition, the HSA modules of SRCAP and TIP60 are oriented oppositely relative to the core module of the respective complexes (Figures 6A and 6B). In addition, while the HSA module is stably anchored to the core of the free TIP60 complex through multivalent contacts with YL1, the SRCAP HSA module is only stabilized upon nucleosome binding, and no cryo-EM density is observed in the structure of the free SRCAP complex (Figures 6A-D).

Comparative structural analyses of HSA modules from human TIP60, human SRCAP, fungal NuA4, and fungal INO80 reveal a conserved architecture (Figures 6E and S10E)^30,42^. However, these modules appear to serve distinct functions across the different complexes. In SRCAP, the HSA module binds to nucleosome DNA at SHL4 to 6, likely facilitating DNA unwrapping for histone exchange (Figure 6E). The INO80 HSA module binds to linker DNA at SHL-9 to -11, functioning to read the linker DNA length and shape (Figure S10E)^42^. In contrast, the TIP60 HSA module, likely devoid of DNA binding capacity due to occlusion by the EPC1/EP400 beta cluster and the C-terminus of EP400, appears to serve an architectural role, as observed for yeast NuA4 (Figures 6E and S10E)^30^.

Comparison of the SRCAP and EP400 ATPase structures revealed striking dissimilarities within their Snf2 family-specific insertions (Figures 6F and 6G). The EP400 ATPase contains a unique ‘latch helix’ that binds the surface of the gating helix that typically inserts into the minor groove upon DNA binding (Figure 6F). Superposition of the nucleosome from the fully engaged SRCAP-nucleosome structure onto the structure of the RUVBL lobe of TIP60 (using the C-lobe of the ATPase for alignment) confirms that the EP400 latch helix would sterically clash with nucleosomal DNA (Figures 6F and 6I). Additionally, the brace motif, which typically bridges the N- and C-lobes in Snf2-family ATPases, adopts an altered conformation in the EP400 ATPase in which it solely interacts with the C-lobe. The lack of a brace between the ATPase lobes allows the N-lobe to form a twisted orientation, resulting in a substantially more open ATPase conformation overall (Figure 6F). This open twisted conformation is stabilized by N-lobe interactions with the EP400 HSA and YL1, and thus significant reorganization of the EP400 ATPase structure would be required for canonical Snf2-family ATPase cleft engagement with DNA (Figures 6D and 6F).

Notably, our ectopically expressed RUVBL lobe cryo-EM sample contained excess nucleosomes at micromolar concentration, yet no nucleosome bound complexes were observed in the cryo-EM data. Moreover, when we measured native SRCAP and TIP60 affinity to 0N0 NCPs, 52N0 nucleosomes, and 147 bp DNA, TIP60 showed almost no binding to NCPs and only weak affinity for linker-containing nucleosomes (52N0) and free DNA (Figure 6H).

In the SRCAP-nucleosome structure, the HSA module forms extensive electrostatic interactions with the nucleosome (Figure 6J). Notably, in the TIP60 nucleosome docked model, the HSA module is in an opposite orientation compared to SRCAP, and there are no nucleosome contacts other than the EP400 ATPase (Figure 6I). While we cannot rule out the possibility of a large conformational change of the HSA module upon nucleosome engagement, it appears to be unlikely as the TIP60 HSA module serves architectural rather than DNA binding functions (Figure 6E). Furthermore, the SRCAP ARP6 anchor domain which recruits the ARP6/ZNHIT1 module is missing from EP400 (Figures 6I, 6J and S10F). As previously detailed, ARP6 and ZNHIT1 subunits in the SRCAP complex unwrap ∼20 bp of nucleosomal DNA near the H2A-H2B dimer that is destined to be evicted (Figures 3F and 6J), and additionally stabilize the nucleosome through ARP6 interactions with the H3 tail and ZNHIT1 binding to the dyad DNA (Figures 3H and 3I). Collectively, the two complexes appear to have evolved divergent structures with associated functional differences.

## Discussion

We present a structural analysis of two paralogous H2A.Z-associated human chromatin remodelers, SRCAP and TIP60. Our comprehensive structural characterization of the human SRCAP complex uncovers a mechanism by which the complex encounters and engages chromatin. The coordinated transition from linker DNA to nucleosome engagement enables SRCAP to utilize 1D diffusion, thereby enhancing target search efficiency within the chromatin environment^31,61–63^. Importantly, the initial interaction with linker DNA would likely drive SRCAP targeting to promoter proximal +1 nucleosomes adjacent to NFRs over gene-body nucleosomes which have shorter linker DNA lengths^10,14^, thus explaining the molecular basis for the enrichment of H2A.Z at +1 nucleosomes observed in vivo^14,17–19^. While our biochemical and structural data underscore the critical role of the nucleosome acidic patch in the nucleosome engagement process, the structural mechanism of ATPase dissociation from the linker DNA and re-association with nucleosomal SHL2 remains unclear. Furthermore, previous work showed the yeast SWR1 complex prefers to exchange histones on canonical H2A-containing nucleosomes with reduced activity on H2A.Z-containing substrates^64,65^. The homologous yeast Swc2 and human YL1 subunits bind to the H2A-H2B acidic patch where H2A and H2A.Z exhibit sequence divergence^66^. Thus, YL1 may also play a role in sensing nucleosome composition prior to engagement.

Most chromatin remodelers across the four main families—SWI/SNF, INO80, ISWI, and CHD—bind to the internal SHL2 site of a nucleosome^24–26,28,29^. Early cryo-EM studies of INO80 suggested that it uniquely binds at SHL-6, but more recent work has shown that it is additionally capable of binding SHL2 on a hexasome substrate^27,67–69^. Remarkably, the nucleosomal conformation of INO80 is reminiscent of the nucleosome-encounter state of SRCAP, and the hexasomal conformation of INO80 is similar to the nucleosome-engaged SRCAP (Figures S7H-K). Biochemical experiments have shown that DNA gaps at SHL2 inhibit INO80 activity on nucleosomes, indicating the nucleosomal SHL-6 conformation may represent an intermediate state prior to full engagement^68,69^. In addition, most chromatin remodelers interact with the acidic patch and can undergo 1D diffusion on free DNA^31,63,70^. Therefore, our proposed structural mechanism of SHL2 engagement through linker DNA and the nucleosome acidic patch may represent a conserved mechanism of nucleosome engagement by other chromatin remodelers.

Consistent with our structure, previous structural and biophysical studies of the yeast SWR1 complex have shown partial DNA unwrapping independent of ATP hydrolysis^22,23,35^. Notably, in contrast to our results and previous SWR1 studies, a recent study reporting multiple cryo-EM structures of the SRCAP-nucleosome complex concluded that ATP hydrolysis is required for partial DNA unwrapping^71^. Mechanistically, we show that the transition from the pre-engaged to nucleosome-engaged state involves a physical wedging of the ZNHIT1-ARP6 subunits between the histone octamer and nucleosomal DNA, resulting in unwrapping of ∼20 bp of DNA. Simultaneously, the ATPase engages the nucleosome and disrupts histone-DNA contacts. In the fully-nucleosome-engaged state, SRCAP is poised for histone exchange. SRCAP envelops the nucleosome and forms extensive interactions likely crucial for subsequent DNA peeling from the histone core and exposure of the H2A-H2B dimer for eviction. In addition, we observe that the H2A.Z-H2B dimer is positioned adjacent to SRCAP ATPase and SRCAP HSA helix, suggesting coordinated histone insertion upon eviction. While our results elucidate the mechanism of nucleosome engagement, how SRCAP exchanges H2A.Z and the role of ATP-hydrolysis remain enigmatic and warrant further investigation.

The TIP60 complex has been implicated in H2A.Z deposition as it shares nine subunits with SRCAP complex^16,20,51–53^. Notably, both complexes natively co-purify with an H2A.Z-H2B dimer and contain paralogous Snf2-type ATPases (EP400/SRCAP). Our comparison of the TIP60 and SRCAP structures reveal critical structural differences despite their shared subunit composition. First, the TIP60-unique latch helix of the EP400 ATPase subunit sterically blocks DNA binding in the cleft, preventing nucleosome engagement and DNA distortion by the gating helix. Second, the TIP60 HSA module is oriented opposite of the nucleosome enveloping SRCAP HSA module, with the ‘A’ submodule of TIP60 serving an architectural rather than DNA binding function as observed in SRCAP. Lastly, TIP60 does not contain the ZNHIT1-ARP6 subunits, which are critical for full nucleosome engagement and DNA unwrapping in the SRCAP complex, and is essential for histone exchange in yeast, flies, mice, and humans^22,72–75^. Collectively, we conclude that the current data do not support a histone exchange function for TIP60.

Early work in Drosophila used overexpressed and affinity purified dPontin (RUVBL1) to obtain dTIP60/Domino complex, which was proposed to have histone exchange activity^51^. Subsequent work, however, showed two isoforms of DOM (Domino) that form distinct complexes with distinct functions^72,76,77^. DOM-A is the functional equivalent of the NuA4/TIP60 complex and displays no H2A.V (Drosophila H2A.Z) exchange activity in vivo, whereas DOM-B is equivalent to the SWR1/SRCAP complex and is required for ATP-dependent H2A.V (H2A.Z) exchange in vivo. Therefore, interpretation of the early work is confounded by the mixture of DOM-A and DOM-B, as well as dINO80 complexes, which also shares the dPontin subunit^51,77^.

Subsequently, studies of human TIP60 showed apparent histone exchange activity in vitro^16,52,53^. However, these studies overexpressed and purified the EP400 subunit in isolation from insect cells^16,52,53^. Our structure shows that EP400 weaves intricately through the RUVBL lobe and interacts with 14 different components of TIP60. EP400 likely does not exist in isolation in cells and is unlikely to be functional in vitro. Moreover, in vivo data shows knock-down of EP400 leads to only minor changes in H2A.Z incorporation^16^. Rather, the TIP60 complex appears to primarily function in acetylating H2A.Z^11,12^. Nevertheless, we cannot rule out the possibility of TIP60 utilizing an entirely different mechanism of histone exchange. In addition, TIP60 may be functional as a chromatin remodeler in unique contexts such as chromatin disrupted by DNA damage, transcription or replication. The physiological roles of TIP60 remain unclear and requires further investigation to unravel.

## Acknowledgements

We thank the JHU Integrated Imaging Center (IIC) for EM instrumentation support. We thank D. Sousa in the Beckman Center for Cryo-EM at Johns Hopkins for microscope access and support. We thank G. McNamara in the High-throughput Phenotypic Screening Core at Johns Hopkins for Tecan SPARK access and support. Cryo-EM image processing was carried out on the Rockfish cluster at the Advanced Research Computing at Hopkins (ARCH) core facility, which is supported by the National Science Foundation (NSF) grant number OAC 1920103. We thank G. Bowman in the Biophysics department at Johns Hopkins for providing acidic patch mutant nucleosomes. This work was funded by the National Institutes of Health (NIH) grants R01GM125831and R35GM149291 to C.W. R.K.L. was supported by NIH Postdoctoral Training Fellowship F32 GM133151.

## Author contributions

R.K.L. conceived of the study, performed cell culturing, generated the gene-edited K-562 cell lines, and constructed recombinant co-expression plasmids. G.P. and R.K.L performed large scale nuclei harvesting for native complex purifications. G.P. purified SRCAP and TIP60 complexes, reconstituted nucleosomes, carried out biochemical assays, and prepared cryo-EM specimens. R.K.L and G.P. collected cryo-EM data. G.P. processed cryo-EM data. A.B.P. built atomic models for the TIP60 complex. R.K.L. built SRCAP models and prepared all final models. G.P. and R.K.L. performed structural analyses. G.P., R.K.L., and A.B.P. prepared figures. G.P. and R.K.L. wrote the paper with input from all authors. R.K.L and C.W. supervised the study.

## Materials and Methods

### Generation of knock-in cell lines

Human K562 cells (ATCC) were cultured at 37 °C and 5% CO2 in RPMI media supplemented with 10% fetal bovine serum and 10 U ml−1 penicillin-streptomycin, and subcultured at a ratio of 1:10 to 1:15 every 2 to 4 d. Wildtype K562 cells were co-transfected with a Cas9 plasmid (CBh-driven 3xFLAG-SV40NLS-hypaSpCas9-NLS; PGK-driven mVenus; U6-driven single-guide (sg) RNA) and a repair plasmid containing Halo-3xFLAG flanked by roughly 800 bp of genomic homology sequence to SRCAP or EPC1 on either side (4.5 μg of repair vector and 1.5 μg of Cas9 vector per T25 flask) using Lipofectamine 3000 (Thermo Fisher) according to the manufacturer’s protocol. Two sgRNAs were designed using the Benchling CRISPR Guide RNA Design Tool (https://www.benchling.com/crispr), cloned into the Cas9 vector and co-transfected with the repair vector individually. After 48 h, transfected cells were combined and sorted using fluorescence activated cell sorting for mVenus fluorescence. mVenus-sorted cells were grown for 7 d, labeled with 50 nM Halo-JF552, and cell populations with higher fluorescence than JF552-labeled wild-type cells were fluorescence activated cell sort-selected and sorted individually into 96-well plates. Clones were expanded and genotyped by PCR. Successfully edited clones were further verified by PCR using multiple primer combinations, Sanger sequencing, and anti-FLAG immunoprecipitation.

### Preparative K562 cell culture and nuclei extraction

Large scale cultures of SRCAP-Halo-3xFLAG and EPC1-Halo-3xFLAG K562 cells were grown at 37 °C and 5% CO2 in Opti-MEM media (Thermo Fisher) supplemented with 0.5% fetal bovine serum and 0.5% newborn calf serum. Cells were maintained in four 3 L spinner flasks (Bellco), each containing 3 L of K562 cultures and constantly stirred at 60 r.p.m. via a Precision Magnetic Stirrer Platform (Bellco). Every 3 or 4 days, cells were split 1:15 or 1:30 into fresh media grown to a density of roughly 7 × 105 cells per ml and collected. To collect, cells were centrifuged using a Fiberlite F9-6×1000 (Thermo Fisher) at 4 °C and 2,500 xg. for 15 min. Cells were washed in PBS, then centrifuged at 4 °C and 2,500 xg for 5 min. Cells were resuspended in 5 volumes of ice cold buffer A (10 mM HEPES pH 7.6, 1.5 mM MgCl2, 10 mM KCl, 1 mM DTT, 1× Roche cOmplete protease inhibitors), immediately pelleted (2,500 xg, 4 °C, 5 min), then resuspended in 3 volumes ice cold buffer A and incubated on ice for 10 minutes^78^. Cells were lysed by douncing ten times using a glass homogenizer with a type B pestle. Nuclei were pelleted by centrifugation (3,000 xg, 4 °C, 20 min), and pellets were flash frozen in liquid nitrogen and stored at −80 °C until use.

### Purification of native complexes

SRCAP and TIP60 were purified similarly. All steps were performed at 4 °C. Frozen nuclei from 50-100 L of cell culture were thawed, 0.9 volumes of buffer C (25 mM HEPES pH 7.6, 0.2 mM EDTA, 25% glycerol, 0.42 M NaCl, 2 mM beta-mercaptoethanol, 0.01% IGEPAL CA-630, 2× Roche cOmplete protease inhibitors, 2× phosphatase inhibitors (1x: 1 mM NaF, 20 mM β-Glycerophosphate)) added and dounced using a glass homogenizer and a type B pestle 20 times on ice. The nuclear extract was nutated for 15 minutes then centrifuged using a JA-20 Beckman rotor at 4 °C and 18,000 xg for 30 min. The supernatant was collected and added to 4 ml of magnetic FLAG M2 resin (Sigma) for 4 h nutating. The resin was washed twice with 10 CV of Column Buffer (25 mM HEPES pH 7.6, 0.3 M NaCl, 1 mM MgCl2, 10% glycerol, 1 mM EDTA, 1 mM beta-mercaptoethanol, 0.01% IGEPAL CA-630, 1× Roche cOmplete protease inhibitors, 1× phosphatase inhibitors), five times with 10 CV of Wash Buffer (Column buffer containing 0.6 M NaCl) and twice with 10 CV of Column Buffer. To elute, the beads were incubated with Column Buffer with 0.5 mg ml−1 3xFLAG peptide rocked for 1 h, then pelleted, and this was repeated for four 1-h elutions. Elutions were concentrated to 300 μl using a 30 kDa molecular weight cutoff Amicon concentrator (Sigma). Concentrated sample was loaded onto a 20%-50% glycerol gradient (25 mM HEPES pH 7.6, 0.3 M NaCl, 1 mM MgCl2, 0.2 mM EDTA, 1 mM DTT, 0.01% IGEPAL CA-630) and ultracentrifuged using a SW55 Ti Beckman rotor at 4 °C and 40,000 rpm for 20 hours. For SRCAP purification, CFDP1 (homolog of yeast Swc5) was bacterially purified and added prior to the glycerol gradient but did not co-migrate with SRCAP complex. Peak fractions of SRCAP/TIP60 complex, assessed by SDS-PAGE stained with Flamingo Fluorescent Stain (Bio-Rad), were pooled and concentrated using a 30 kDa molecular weight cutoff Amicon concentrator (Sigma). The concentrated sample was used immediately for cryo-EM or frozen in liquid nitrogen and stored at −80 °C.

### RUVBL lobe expression and purification

The 8 open reading frames of the RUVBL lobe (RUVBL1, RUVBL2, YL1, EP400, EPC1, GAS41, BAF53a, DMAP1) were sub-cloned into modified pCAG vectors. The P2A self-cleaving peptide coding sequence was inserted between RUVBL1 and RUVBL2. All other subunits were driven by individual CAG promoters. To isolate the RUVBL lobe, residues 1-657 of EP400 (EP400ΔN) and residues 1-358 and 621-836 of EPC1 were deleted. For protein purification, RUVBL2 was MBP tagged and EP400ΔN was HaloTag and 3xFLAG tagged. Expi293 cells were grown in Expi293 Expression medium (Gibco) to a density of roughly 2 × 106 cells per ml at 37 °C and 8% CO2. The plasmids were co-transfected using polyethylenimine (1 mg total DNA and 3 mg PEI per 1 L culture). The transfected cells were cultured for 72 hours at 37 °C and 8% CO2 and harvested by centrifugation. All purification steps were performed at 4 °C. Cells were resuspended in Lysis Buffer (50 mM HEPES pH 7.6, 0.4 M NaCl, 0.25% CHAPS, 5 mM ATP, 5 mM MgCl2, 0.25 mM EDTA, 2 mM DTT, 10% glycerol, 1× Roche cOmplete protease inhibitors, 1× phosphatase inhibitors) and dounced using a glass homogenizer and a type B pestle 20 times on ice. The nuclear extract was nutated for 30 minutes then centrifuged using a JA-20 Beckman rotor at 4 °C and 22,000 xg for 30 min. The supernatant was collected and incubated with 10 ml of Amylose resin (New England Biolabs) for 90 minutes. The resin was washed five times with 10 CV of Amylose Buffer (30 mM HEPES pH 7.6, 0.3 M NaCl, 0.1% CHAPS, 0.25 mM EDTA, 2 mM DTT, 10% glycerol, 1× Roche cOmplete protease inhibitors). Protein was eluted with Amylose Buffer supplemented with 20 mM maltose. The amylose elution was then transferred to 2 ml of magnetic FLAG M2 resin (Sigma) and incubated for 3 hours. The FLAG resin was washed and eluted similarly as described for the native TIP60 complex. The FLAG elutions were concentrated to 300 μl using a 30 kDa molecular weight cutoff Amicon concentrator (Sigma). Concentrated sample was loaded onto a 15%-45% glycerol gradient (25 mM HEPES pH 7.6, 0.25 M NaCl, 0.2 mM EDTA, 1 mM DTT, 0.01% IGEPAL CA-630) and ultracentrifuged using a SW55 Ti Beckman rotor at 4 °C and 37,000 rpm for 17 hours. Peak fractions, assessed by SDS-PAGE stained with Flamingo Fluorescent Stain (Bio-Rad), were pooled and concentrated using a 30 kDa molecular weight cutoff Amicon concentrator (Sigma). The concentrated sample was used immediately for cryo-EM or frozen in liquid nitrogen and stored at −80 °C.

### TRRAP lobe expression and purification

The 6 open reading frames of the TRRAP lobe (TRRAP, EP400, EPC1, GAS41, BAF53a, DMAP1) were sub-cloned into modified pCAG vectors. All subunits were driven by individual CAG promoters. To isolate the TRRAP lobe, residues 1-512, 914-2132, and 2562-3159 of EP400 (EP400ΔM) and residues 1-304 of EPC1 were deleted. For protein purification, EP400ΔM was HaloTag and 6xHIS tagged and TRRAP was 2xFLAG tagged. Expi293 cells were transfected, cultured, harvested, and lysed similarly as described for the RUVBL lobe. The clarified lysate was incubated with 2 ml of magnetic FLAG M2 resin (Sigma) and incubated for 3 hours. The FLAG resin was washed and eluted similarly as described for the native TIP60 complex, except for the omission of EDTA from the elution buffer. The FLAG elution was transferred to 2 ml of NI-NTA agarose resin (Qiagen) and incubated for 1 hour. The resin was washed five times with 10 CV of Nickel Buffer (25 mM HEPES pH 7.6, 0.3 M NaCl, 10 mM imidazole, 2 mM DTT, 0.01% IGEPAL CA-630, 10% glycerol, 1× Roche cOmplete protease inhibitors). Protein was eluted with Nickel Buffer supplemented with 300 mM imidazole. The NI-NTA elution was concentrated and loaded onto a glycerol gradient similarly as for the RUVBL lobe. The sample was used immediately for cryo-EM or frozen in liquid nitrogen and stored at −80 °C.

### Preparation of nucleosome substrates

Recombinant Xenopus histones H2A, H2B, H3, and H4 were expressed and purified as described previously^79^. The nucleosomal DNA contained the upstream super core promoter^80^ and downstream TERT promoter followed by a modified Widom 601 sequence^81^ and random linker sequence. The 285-bp DNA sequence was 5’-ATCGAAGGGCGCCTATATAAGGGGGTGGGGGCGCGTTCGTCCTCCC TCTCCTCGCGGCGCGAGTTTCAGGCAGCGCTGCGTCCTGCTGCGCA CGTGGGAAGCCCTG<u>CTGGAGAATCCCGGTGCGCAGGCCGCTCAATT GGTCGTAGACAGCTCTAGCACCGCTTAAACGCAGCTACGCGCTGTC CCCCGCGTTTTAACCGCCAAGGGGATTACTCCCTAGTCTCCAGGCA GCTGTCAGATATGTACATCCTGT</u>GATCCCCGGGTACCGAGCTCGAA TTCACTGGC - 3’. The Widom 601 sequence is underlined. The DNA was generated by large scale polymerase chain reaction, purified by anion exchange chromatography and ethanol precipitation. For nucleosome reconstitution, DNA and histone octamers were mixed in equimolar ratio and dialyzed against a gradient of decreasing salt concentration^79^. Reconstituted nucleosomes were purified on a Model 491 Prep Cell (Bio-Rad), concentrated to 5 uM and stored at 4 °C. The acidic patch mutant (APM) nucleosomes were provided by G. Bowman and generated similarly but with the following 225-bp DNA sequence: 5’-<u>TGGAGAATCCCGGTGCCGAGGCCGCTCAATTGGTCGTAGACAGCTC TAGCACCGCTTAAACGCACGTACGCGCTGTCCCCCGCGTTTTAACC GCCAAGGGGATTACTCCCTAGTCTCCAGGCACGTGTCAGATATATA CATCCTG</u>TGCATGTATTGAACAGCGACCTTGCCGGTGCCAGTCGGA TAGTGTTCCGAGCTCCCACTCTAGAGGATCCCCGGGTACCG - 3’. The substrates used for AlphaScreen (WT NCP, APM NCP, WT 52N0, 147 bp DNA, 199 bp DNA) were purchased from Epicypher (dCypherTM Nucleosome Full Panel).

### Nucleosome binding assays

The dCypher-binding assay to nucleosomes were performed as previously ^30^. Briefly, 2.5 ul of native SRCAP (25 nM final) was incubated with 2.5 ul of nucleosome (10 nM final) for 30 min at room temperature in binding buffer (25 mM HEPES pH 7.6, 0.25 mM EGTA, 0.25 mM EDTA, 2.5 mM MgCl2, 0.25 mg/ml BSA, 70 mM NaCl, 5% glycerol, 0.025% IGEPAL CA-630, 1 mM DTT). Then 2.5 ul of 0.1 mg/ml AlphaScreen Streptavidin Donor Beads (6760003) and 2.5 ul of 0.1 mg/ml AlphaScreen anti-FLAG A Acceptor Beads (6760128) were added, followed by a 45-min incubation at room temperature. Alpha counts were measured on a Tecan SPARK microplate reader (150 ms excitation time, 680 nM laser excitation, 570 nm emission filter ± 50 nm bandwidth). Binding assays were performed in triplicates and the mean and standard deviation are shown. For the EMSA, Cy5 labeled WT and acidic patch mutant (APM) nucleosomes (0N80, 5 nM) was incubated with native SRCAP (0, 5, 10, 25, 50 nM) for 15 minutes at 37 C in binding buffer (supplemented with 1 mM ATPyS). Then the reaction was resolved on a 0.8% agarose gel and scanned on a Typhoon biomolecular imager (Cytiva). Binding assays were performed in triplicates and the mean and standard deviation are shown.

### Negative stain EM sample preparation, data collection, and image processing

3 μl of native TIP60 complex was diluted to 100 nM in NS-EM buffer (25 mM HEPES pH 7.6, 2.5 mM MgCl2, 0.25 M NaCl, 2 mM DTT, 0.01% IGEPAL CA-530) and absorbed to glow discharged (Tergeo-EM, PIE Scientific) continuous carbon grids for 5 minutes and stained with 1% uranyl formate. Negative stain data were collected on a Thermo Fisher Talos F200C at 200 keV equipped with a CETA camera. 1,050 micrographs were collected at a magnification of 57,000 (2.63 Å/pixel) at a defocus range of -1.0 to -1.5 μm using EPU software (Thermo Fisher). CryoSPARC was used for picking and extracting particles, 2D classification, and 3D refinement^82^.

### Cryo-EM sample preparation

#### Native SRCAP

For the SRCAP-ATPyS-106N32 sample (data set 1), native SRCAP (1.4 uM) was incubated with 106N32 nucleosomes (1.8 uM) and ATPyS (1 mM) for 10 minutes at RT then transferred to ice. The final buffer was 25 mM HEPES pH 7.6, 0.2 mM EDTA, 1 mM MgCl2, 85 mM NaCl, 0.25 mM TCEP. 3 ul of the sample was crosslinked with 0.05% glutaraldehyde (Electron Microscopy Sciences) for 3 minutes on ice then applied to glow discharged (Tergeo-EM, PIE Scientific) Quantifoil 0.6/1.0 grids at 4 °C under 100% humidity. The sample was immediately blotted for 5 s at 5 N force and vitrified in liquid ethane using Vitrobot Mark IV (FEI). For the SRCAPATPyS-0N80 (APM) sample (data set 2), native SRCAP (1.0 uM) was incubated with 0N80 (APM) nucleosomes (1.5 uM) and ATPyS (1 mM) for 30 minutes at RT then transferred to ice. The final buffer, glow discharging, crosslinking, and blotting conditions were identical as data set 1 except that Quantifoil 1.2/1.3 grids were used. For the SRCAP-ATPyS-106N32 sample (data set 3), native SRCAP (1.0 uM) was incubated with 106N32 nucleosomes (1.7 uM) and ATPyS (1 mM) for 30 minutes at RT, 3 minutes at 37 C, then transferred to ice. The final buffer, glow discharging, crosslinking, and blotting conditions were identical as data set 1. The SRCAP-106N32 (no nucleotide) sample (data set 4) was prepared identically as data set 3 except no ATPyS was added.

#### Native TIP60

3 μl of diluted native TIP60 complex (0.3 uM) in TIP60 buffer (25 mM HEPES pH 7.6, 0.2 mM EDTA, 2.5 mM MgCl2, 0.25 M NaCl, 2 mM DTT, 0.01% IGEPAL CA-530, 3% glycerol) was mixed with 0.3 ul of 0.5% glutaraldehyde (0.05% final, Electron Microscopy Sciences) and immediately applied to a graphene oxide coated Holey Carbon 2/1 grids (Quantifoil) ^83^. The sample was incubated for 5 minutes at 4 °C under 100% humidity then blotted for 5 s at 0 N force and vitrified in liquid ethane using Vitrobot Mark IV (FEI).

#### RUVBL lobe

The RUVBL lobe (1.0 uM) was incubated with ATPyS (1 mM) for 5 minutes at RT, and transferred on ice. The final buffer was 25 mM HEPES pH 7.6, 0.2 mM EDTA, 1 mM MgCl2, 0.1 M NaCl, 1 mM DTT, 0.25 mM TCEP, 0.001% IGEPAL CA-530, 1% glycerol. 3 μl of sample was crosslinked with 0.05% glutaraldehyde (Electron Microscopy Sciences) for 5 minutes on ice then applied to glow discharged (Tergeo-EM, PIE Scientific) Au-FLAT 1.2/1.3 grids (Protochip) at 4 °C under 100% humidity. The sample was immediately blotted for 4 s at 1 N force and vitrified in liquid ethane using Vitrobot Mark IV (FEI).

#### TRRAP lobe

3 μl of TRRAP lobe (1.0 uM) in TRRAP buffer (25 mM HEPES pH 7.6, 0.2 mM EDTA, 1 mM MgCl2, 0.25 M NaCl, 2 mM DTT, 0.005% IGEPAL CA-530, 1% glycerol) was crosslinked with 0.05% glutaraldehyde (Electron Microscopy Sciences) for 5 minutes on ice then applied to glow discharged (Tergeo-EM, PIE Scientific) Au-FLAT 1.2/1.3 grids (Protochip) at 4 °C under 100% humidity. The sample was immediately blotted for 4 s at 1 N force and vitrified in liquid ethane using Vitrobot Mark IV (FEI).

### Cryo-EM data collection

#### SRCAP

Data set 1 was collected on a Thermo Fisher G3i Titan Krios at 300 keV equipped with a K3 direct electron-counting camera (Gatan). Semi-automated data collection was done using the SerialEM data collection software^84^. For data set 1, 20,709 micrographs were collected at a magnification of 22,500 (1.025 Å/pixel, super-resolution mode) at a defocus range of -0.8 to -1.6 μm. Each micrograph was 64 frames with 4.0 total exposure time and a total electron dose of ∼50 e−/Å^2^. Data sets 2-4 were collected on a Thermo Fisher G3i Titan Krios at 300 keV equipped with a Falcon 4i direct electron detector (Thermo Fisher) and Selectris Energy Filter (Thermo Fisher). Automated data collection was done using the EPU software (Thermo Fisher). For data set 2, 9,744 micrographs were collected at a magnification of 165,000 (0.726 Å/pixel) at a defocus range of -0.8 to - 1.6 μm. Each micrograph was 1,197 frames with 3.89 s total exposure time and a total electron dose of ∼40 e−/Å^2^. For data set 3 and 4, 6,417 and 6,724 micrographs were collected at a magnification of 165,000 (0.726 Å/pixel) at a defocus range of -0.8 to -1.6 μm. Each micrograph was 873 frames with 2.84 s total exposure time and a total electron dose of ∼40 e−/Å^2^.

#### TIP60

Cryo-EM data of native TIP60, RUVBL lobe, and TRRAP lobe were collected on a Thermo Fisher G3i Titan Krios at 300 keV equipped with a Falcon4i direct electron detector (Thermo Fisher) and Selectris Energy Filter (Thermo Fisher). Automated data collection was done using the EPU software (Thermo Fisher). For the native TIP60 complex, 28,451 micrographs were collected at a magnification of 130,000 (0.925 Å/pixel) at a defocus range of -1.0 to -2.5 μm. Each micrograph was 1,134 frames with 3.69 s total exposure time and a total electron dose of ∼40 e−/Å^2^. For the RUVBL lobe, 13,480 total micrographs (8229, 5251) were collected at a magnification of 165,000 (0.726 Å/pixel) at a defocus range of -0.8 to -1.6 μm in two separate sessions from the same grid. Each micrograph was 1,113 frames with 4.62 s total exposure time and a total electron dose of ∼40 e−/Å^2^. For the TRRAP lobe, 12,277 micrographs were collected at a magnification of 165,000 (0.726 Å/pixel) at a defocus range of -0.8 to -1.6 μm. Each micrograph was 954 frames with 3.10 s total exposure time and a total electron dose of ∼40 e−/Å^2^.

### SRCAP Cryo-EM image processing

Cryo-EM data was processed using Relion v4.0^85^. Movie frames were aligned using MotionCor2 (data set 1) or Relion’s own implementation (data set 2-4) to correct for specimen motion^85,86^. CTF parameters were estimated using Gctf (data set 1) or CTFFIND4 (data set 2-4)^87,88^. For data set 1, 7,394,086 particles were picked with the LoG picker and 10,833,376 particles were picked using 2D class averages of free SRCAP. Both sets of particles were extracted and sorted extensively through multiple rounds of 3D classification (Figures S1 and S2). Through various sorting methods, six states (fully-engaged, partially-engaged, pre-engaged, nucleosome-encounter, free, dimer) were determined from data set 1 (Figures S1, S2, S4, and S5). 227,030 fully-engaged particles were 3D refined to 3.35 Å and focused refinements of the core, ARP6/ZNHIT1, ATPase, and nucleosome yielded resolutions of 2.60 Å, 3.16 Å, 3.22 Å, and 3.04 Å. The trident submodule, A submodule, and GAS41 submodule were 3D refined from a subset of particles to 3.84 Å, 9.65 Å, and 7.32 Å. 24,701 partially-engaged particles were 3D refined to 3.92 Å and focused refinements of the core, ARP6/ZNHIT1, and ATPase-nucleosome yielded resolutions of 3.53 Å, 3.53 Å, and 3.88 Å. 28,381 pre-engaged particles were 3D refined to 4.04 Å and focused refinements of the core, ARP6/ZNHIT1, HSA module, and nucleosome yielded resolutions of 3.66 Å, 3.96 Å, 7.61 Å, and 5.34 Å. 18,928 nucleosome-encounter particles were 3D refined to 4.30 Å and focused refinements of the core, ARP6/ZNHIT1, ATPase, and nucleosome yielded resolutions of 3.53 Å, 3.92 Å, 4.35 Å, 7.91 Å. 37,293 free SRCAP particles were 3D refined to 3.56 Å and focused refinements of the core and ARP6/ZNHIT1 yielded resolutions of 3.41 Å and 3.66 Å. 8,397 dimer particles were 3D refined to 6.35 Å and focused refinements of copy 1 and copy 2 yielded resolutions of 4.32 Å and 4.09 Å. For data set 2, 2,046,788 particles were picked with the LoG picker and 478,824 particles were picked using 2D class averages of free SRCAP. Both sets of particles were extracted and sorted extensively through multiple rounds of 3D classification (Figures S3B). Only the linker DNA-bound and nucleosome-encounter (APM) states were observed. 31,816 linker DNA-bound particles were 3D refined to 3.87 Å and focused refinements of core, ARP6/ZNHIT1, and ATPase yielded resolutions of 3.70 Å, 3.62 Å, and 4.01 Å. 42,861 nucleosome-encounter (APM) particles were 3D refined to 3.67 Å and focused refinements of core, ARP6/ZNHIT1, ATPase, and nucleosome yielded resolutions of 3.40 Å, 3.55 Å, 4.09 Å, and 10.13 Å. For data set 3, 718,381 particles were picked with the LoG picker and 981,281 particles were picked using 2D class averages of free SRCAP. Both sets of particles were extracted and sorted extensively through multiple rounds of 3D classification (Figure S3D). 12,372 nucleosome-encounter (WT) particles were 3D refined to 3.94 Å and focused refinements of core, ARP6/ZNHIT1, ATPase, and nucleosome yielded resolutions of 3.50 Å, 3.69 Å, 4.22 Å, and 5.08 Å. 13,297 particles were used to generate the fully-engaged state which was not analyzed further. For data set 4, 791,566 particles were picked with the LoG picker and 1,051,835 particles were picked using 2D class averages of free SRCAP. Both sets of particles were extracted and sorted extensively through multiple rounds of 3D classification (Figure S3F). 4,625 particles were used to generate the pre-engaged state and 5,947 particles were used for the nucleosome-encounter state, both of which were not further analyzed. All resolutions reported were determined from gold-standard Fourier shell correlation (FSC) of 0.143^89^.

### TIP60 Cryo-EM Image processing

#### Native TIP60

Cryo-EM data was initially processed with CryoSPARC v4.2.1 to template pick particles, then transferred for further processing on Relion v4.0^82,85^. Movie frames were aligned to correct for specimen motion. The CTF parameters were estimated using CTFFIND4^88^. Particles were extracted in Relion from coordinates obtained with CryoSPARC for the RUVBL lobe and the TRRAP lobe. For the RUVBL lobe, 1,762,267 coordinates were extracted and binned by 4. Two rounds of 3D classification resulted in 130,599 particles. Selected particles were subjected to 3D refinement and re-extracted and binned by 2. One round of 3D classification resulted in 77,683 particles. Selected particles were subjected to 3D refinement and re-extracted with no binning. After one round of CTF refinement, particle polishing, and 3D refinement, another round of 3D classification was performed 90,91. 61,163 particles were used for the final 3D refinement which resulted in a gold standard resolution of 3.43 Å. To improve the map quality of the flexible portions, focus refinement was performed and the focused maps were combined into a composite map. The resolution of the bottom RUVBL, top RUVBL, and HSA module were 3.13 Å, 3.57 Å, 3.40 Å, respectively. All resolutions reported were determined from gold-standard Fourier shell correlation (FSC) of 0.143. For the TRRAP lobe, 2,448,844 coordinates were extracted and binned by 4. One round of 3D classification resulted in 1,406,651 particles. Selected particles were subjected to 3D refinement and re-extracted and binned by 2. One round of 3D classification resulted in 337,773 particles. Selected particles were subjected to 3D refinement and re-extracted with no binning. Another round of 3D classification resulted in 106,666 particles. Selected particles were subjected to one round of CTF refinement, and particle polishing. The 106,666 particles were used for the final 3D refinement which resulted in a gold standard resolution of 3.05 Å. To improve the map quality of the flexible portions, focus refinement was performed and the focused maps were combined into a composite map. The resolution of the Core, Top, Bottom, and Distal were 2.81 Å, 3.08 Å, 3.25 Å, 3.43 Å, respectively. All resolutions reported were determined from gold-standard Fourier shell correlation (FSC) of 0.143^89^.

#### RUVBL lobe

Cryo-EM data was processed using Relion v4.0^85^. The two datasets were processed independently then later combined for refinements. Movie frames were aligned and CTF parameters were estimated similarly as in the native TIP60 sample^88^. 765,637 and 397,577 particles were picked using Relion LoG picker and extracted. The two sets of particles were independently subjected to one round of 2D classification, and two rounds of 3D classification which resulted in 22,482 and 26,503 particles. The particles were subjected to one round of CTF refinement, particle polishing, and 3D refinement 90,91. The refined particles were combined for a total of 45,284 particles. The 45,284 particles were used for the final 3D refinement which resulted in a gold standard resolution of 3.07 Å. To improve the map quality of the flexible portions, focus refinement was performed and the focused maps were combined into a composite map. The resolution of the bottom RUVBL, top RUVBL, HSA module, and ATPase were 2.72 Å, 3.54 Å, 3.02 Å, 3.58 Å, respectively. All resolutions reported were determined from gold-standard Fourier shell correlation (FSC) of 0.143^89^.

#### TRRAP lobe

Cryo-EM data was processed similarly as the native TIP60. Briefly, CryoSPARC was used to template pick 695,891 particles then the coordinates were used to extract the particles in Relion v4.0^82,85^. Movie frames were aligned and CTF parameters were estimated similarly as in the native TIP60 sample^88^. Three rounds of 3D classification resulted in 141,352 particles which were subjected to two rounds of CTF refinement, particle polishing, and 3D refinement 90,91. The 141,352 particles were used for the final 3D refinement which resulted in a gold standard resolution of 2.35 Å. To improve the map quality of the flexible portions, focus refinement was performed and the focused maps were combined into a composite map. The resolution of the Core, Top, Bottom, and Distal were 2.25 Å, 2.65 Å, 2.88 Å, 2.97 Å, respectively. All resolutions reported were determined from gold-standard Fourier shell correlation (FSC) of 0.14389.

### Model building and refinement

In order to construct comprehensive atomic models for the SRCAP and TIP60 structures, the focused-refined cryo-EM maps were post-processed using DeepEMhancer^92^ and then combined into composite maps after aligning them to one another using the globally refined structure as a reference, and removing overlapping densities via tight masking. Additional maps, e.g. those post-processed by local-resolution filtering, were also used to guide model construction in some cases. The refined models deposited to the PDB from this study were generally restricted to regions resolved better than 5 Å resolution in the cryo-EM maps. Initial models were largely constructed from Alphafold2-predicted structures. Manual model building was performed in Coot^93^ followed by flexible fitting using ISOLDE in Chimera X^94,95^. Models were then subjected to iterative cycles of real-space refinement in PHENIX^96^ and manual outlier adjustment in ISOLDE. The final models were validated using MTriage^97^ and MolProbity^98^ within PHENIX. Extended models were also constructed which include additional components beyond the refined PDB-deposited models, in instances where structures could be reliably fit into lower resolution densities (generally between 5-20 Å) in the cryo-EM maps. These extended models are provided as Supplementary Data. For the SRCAP structures, the fully nucleosome-engaged state exhibited the highest resolution and completeness, and thus this structure was built and refined first. The fully engaged SRCAP-nucleosome model was then used to generate starting models for the remaining SRCAP structures, and starting model restraints were enforced within PHENIX and ISOLDE to limit divergence from the starting model. Similarly, the models for the TIP60 RUVBL and TRRAP lobes were first built into the higher resolution reconstructions obtained from the ectopically expressed complexes, and these were used as starting models for the structures obtained from the endogenously purified native TIP60 complex. AlphaFold2 predictions were carried out using the ColabFold AlphaFold2_mmseqs2 Notebook on GoogleColab (https://colab.research.google.com/github/sokrypton/ColabFold/blob/main/AlphaFold2.ipynb)^56–58^.

### Figure generation

Figures were generated using UCSF ChimeraX^95,99^, Adobe Illustrator, and GraphPad Prism.

### Statistics and reproducibility

Statistical analyses were performed using GraphPad Prism (v10.2.3). All biochemical experiments were performed in triplicates and values reported are mean ±standard deviation. Significance levels: *P ≤ 0.05, **P ≤ 0.01, and ***P ≤ 0.001. Statistical tests and P values for figure 6H are reported in the figure legends.

**Table S1:** Cryo-EM data collection, refinement and validation statistics for SRCAP structures.

|  | SRCAP-nucleosome, fully engaged (EMDB-45381) (PDB 9CA7) | SRCAP-nucleosome, partially engaged (EMDB-45382) (PDB 9CA8) | SRCAP-nucleosome, pre-engaged (EMDB-45384) (PDB 9CAA) | Unbound SRCAP (EMDB-45383) (PDB 9CA9) | SRCAP-nucleosome encounter (EMDB-45385) (PDB 9CAB) |
| --- | --- | --- | --- | --- | --- |
| **Data collection and processing** |  |  |  |  |  |
| Data set | Data set 1 | Data set 1 | Data set 1 | Data set 1 | Data set 3 |
| Microscope | Krios G3i | Krios G3i | Krios G3i | Krios G3i | Krios G3i |
| Voltage (kV) | 300 | 300 | 300 | 300 | 300 |
| Nominal magnification | 22,500 | 22,500 | 22,500 | 22,500 | 165,000 |
| Energy filter | none | none | none | none | Selectris |
| Energy slit width |  |  |  |  | 10 eV |
| Detector | K3 | K3 | K3 | K3 | Falcon 4i |
| Pixel size (Å) | 1.025 | 1.025 | 1.025 | 1.025 | 0.726 |
| Defocus range (µm) | -0.8 to -1.6 | -0.8 to -1.6 | -0.8 to -1.6 | -0.8 to -1.6 | -0.8 to -1.6 |
| Exposure time | 4.0 | 4.0 | 4.0 | 4.0 | 2.8 |
| Electron exposure (e<sup>-</sup>/Å<sup>2</sup>) | 50 | 50 | 50 | 50 | 40 |
| Movies collected | 20,709 | 20,709 | 20,709 | 20,709 | 6,417 |
| **Cryo-EM Reconstruction** |  |  |  |  |  |
| Software | RELION | RELION | RELION | RELION | RELION |
| Particle images (no.) | 227,030 | 24,701 | 28,381 | 37,293 | 12,372 |
| Consensus map resolution, FSC = 0.143 (Å) | 3.35 | 3.92 | 4.04 | 3.56 | 3.94 |
| **Coordinates Refinement** |  |  |  |  |  |
| Software | PHENIX | PHENIX | PHENIX | PHENIX | PHENIX |
| Map vs. model resolution, FSC = 0.5 (Å) | 2.6 | 3.4 | 3.9 | 3.2 | 4.0 |
| Map vs. model cross-correlation | 0.85 | 0.76 | 0.67 | 0.79 | 0.66 |
| Model composition |  |  |  |  |  |
| Non-hydrogen atoms | 42370 | 42954 | 41005 | 28594 | 44309 |
| Protein residues | 4709 | 4709 | 4386 | 3618 | 4700 |
| DNA residues | 242 | 270 | 306 | 0 | 338 |
| Ligands | AGS(2), ADP(6)<br>Mg<sup>2+</sup>(5), Zn<sup>2+</sup>(2) | AGS(2), ADP(6)<br>Mg<sup>2+</sup>(5), Zn<sup>2+</sup>(2) | AGS(1), ADP(6)<br>Mg<sup>2+</sup>(4), Zn<sup>2+</sup>(2) | AGS(1), ADP(6)<br>Mg<sup>2+</sup>(4), Zn<sup>2+</sup>(2) | AGS(2), ADP(6)<br>Mg<sup>2+</sup>(5), Zn<sup>2+</sup>(2) |
| Average B factors (Å<sup>2</sup>) |  |  |  |  |  |
| Protein | 59 | 95 | 108 | 97 | 116 |
| DNA | 71 | 140 | 135 |  | 165 |
| Ligand | 48 | 88 | 89 | 72 | 92 |
| R.m.s. deviations |  |  |  |  |  |
| Bond lengths (Å) | 0.004 | 0.003 | 0.002 | 0.003 | 0.002 |
| Bond angles (°) | 0.554 | 0.509 | 0.489 | 0.482 | 0.510 |
| Validation |  |  |  |  |  |
| MolProbity score | 1.21 | 1.18 | 1.19 | 1.20 | 1.15 |
| All-atom clashscore | 4.27 | 3.95 | 4.06 | 4.12 | 3.56 |
| Poor rotamers (%) | 0 | 0 | 0 | 0 | 0 |
| Ramachandran plot |  |  |  |  |  |
| Favored (%) | 98.1 | 98.2 | 98.1 | 98.0 | 98.3 |
| Allowed (%) | 1.9 | 1.8 | 1.9 | 2.0 | 1.7 |
| Disallowed (%) | 0.0 | 0.0 | 0.0 | 0.0 | 0.0 |

**Table S2:** Cryo-EM data collection, refinement and validation statistics for TIP60 structures.

|  | Native TIP60<br>TRRAP module<br>(EMDB-45387)<br>(PDB 9CAD) | Native TIP60<br>RuvBL module<br>(EMDB-45386)<br>(PDB 9CAC) | Recombinant<br>TRRAP module<br>(EMDB-45389)<br>(PDB 9CAF) | Recombinant<br>RuvBL module<br>(EMDB-45388)<br>(PDB 9CAE) |
| --- | --- | --- | --- | --- |
| **Data collection and processing** |  |  |  |  |
| Microscope | Krios G3i | Krios G3i | Krios G3i | Krios G3i |
| Voltage (kV) | 300 | 300 | 300 | 300 |
| Magnification | 130,000 | 165,000 | 165,000 | 165,000 |
| Energy filter | Selectris | Selectris | Selectris | Selectris |
| Energy slit width | 10 eV | 10 eV | 10 eV | 10 eV |
| Detector | Falcon 4i | Falcon 4i | Falcon 4i | Falcon 4i |
| Pixel size (Å) | 0.925 | 0.925 | 0.726 | 0.726 |
| Defocus range (µm) | -1.0 to -2.5 | -1.0 to -2.5 | -0.8 to -1.6 | -0.8 to -1.6 |
| Exposure time | 3.69 | 3.69 | 3.10 | 4.62 |
| Electron exposure (e-/Å<sup>2</sup>) | 40 | 40 | 40 | 40 |
| Movies collected | 28,451 | 28,451 | 12,277 | 13,480 |
| **Cryo-EM Reconstruction** |  |  |  |  |
| Software | RELION | RELION | RELION | RELION |
| Particle images (no.) | 106,666 | 61,163 | 141,352 | 45,284 |
| Consensus map resolution,<br>FSC = 0.143 (Å) | 3.05 | 3.43 | 2.35 | 3.07 |
| **Coordinates Refinement** |  |  |  |  |
| Software | PHENIX | PHENIX | PHENIX | PHENIX |
| Map vs. model resolution,<br>FSC = 0.5 (Å) | 2.7 | 3.0 | 2.0 | 2.4 |
| Map vs. model<br>cross-correlation | 0.80 | 0.73 | 0.84 | 0.77 |
| Model composition |  |  |  |  |
| Non-hydrogen atoms | 29915 | 40861 | 29921 | 33978 |
| Protein residues | 3723 | 5146 | 3724 | 4289 |
| Ligands | IHP (1) | ATP (1), ADP (7),<br>MG (8) | IHP (1) | AGS (8), MG (8) |
| **Average B factors (Å<sup>2</sup>)** |  |  |  |  |
| Protein | 63 | 97 | 39 | 59 |
| Ligand | 81 | 62 | 38 | 38 |
| **R.m.s. deviations** |  |  |  |  |
| Bond lengths (Å) | 0.003 | 0.003 | 0.004 | 0.003 |
| Bond angles (°) | 0.543 | 0.533 | 0.592 | 0.547 |
| **Validation** |  |  |  |  |
| MolProbity score | 1.12 | 1.16 | 1.06 | 1.09 |
| All-atom clashscore | 3.30 | 3.69 | 2.72 | 2.97 |
| Poor rotamers (%) | 0.0 | 0.0 | 0.0 | 0.0 |
| **Ramachandran plot** |  |  |  |  |
| Favored (%) | 98.4 | 98.4 | 98.4 | 98.4 |
| Allowed (%) | 1.6 | 1.6 | 1.6 | 1.6 |
| Disallowed (%) | 0.0 | 0.0 | 0.0 | 0.0 |

**Figure S1:**
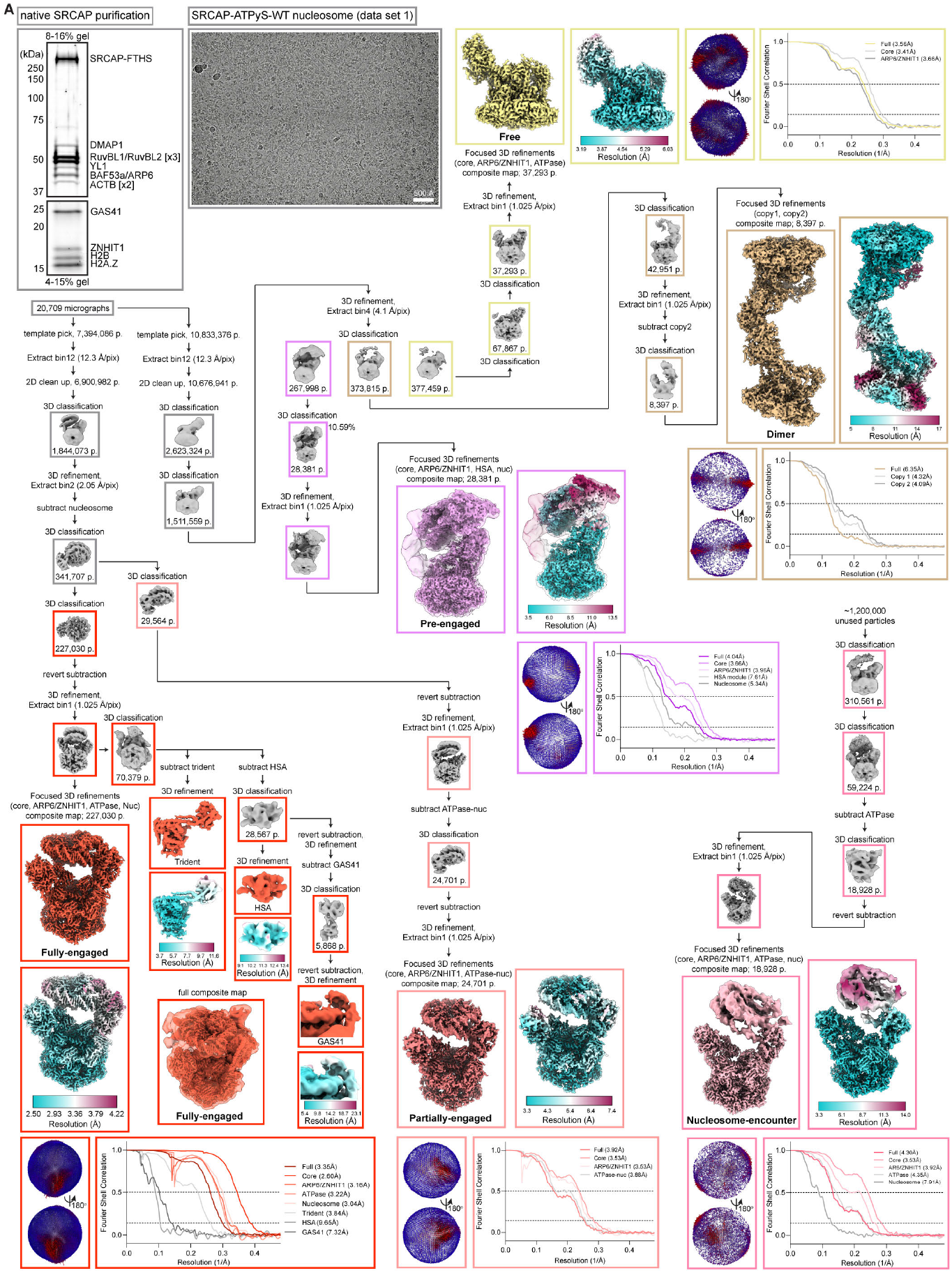
Cryo-EM processing of SRCAP (data set 1). **(A)** SDS-PAGE gel, cryo-EM micrograph, and cryo-EM data processing (data set 1) of SRCAP-ATPγS-WT nucleosome. The post processed composite map, local resolution colored map, mask corrected gold-standard Fourier shell correlation (FSC) plots, and orientation distribution plots for global refinements are shown for each state.

**Figure S2:**
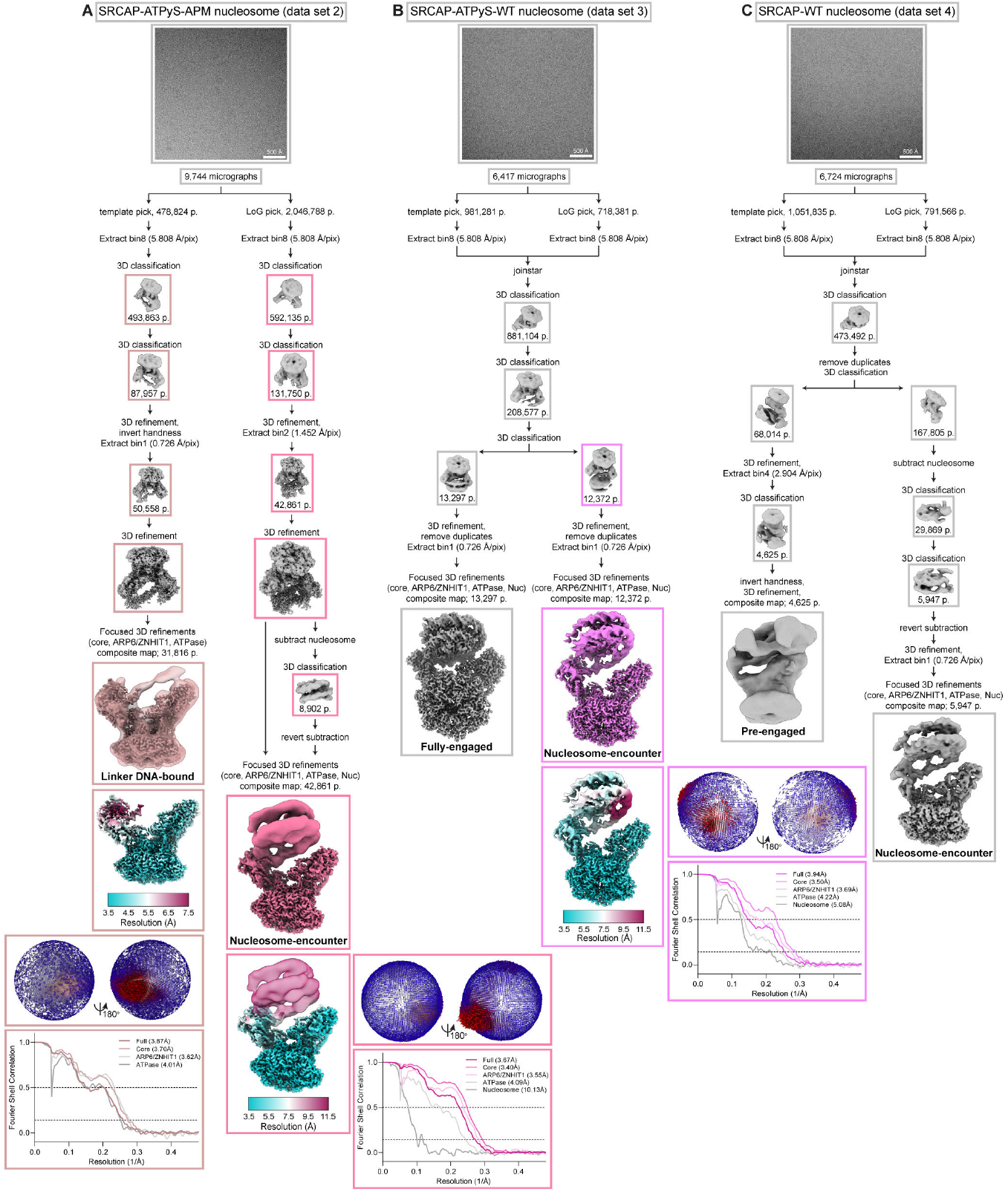
Cryo-EM processing of SRCAP (data set 2-4). **(A-C)** Cryo-EM micrograph and cryo-EM data processing SRCAP-ATPγS-APM nucleosome (A, data set 2), SRCAP-ATPγS-WT nucleosome (B, data set 3), and SRCAP-WT nucleosome (C, data set 4). The post processed composite map, local resolution colored map, mask corrected gold-standard Fourier shell correlation (FSC) plots, and orientation distribution plots for global refinements are shown for each state.

**Figure S3:**
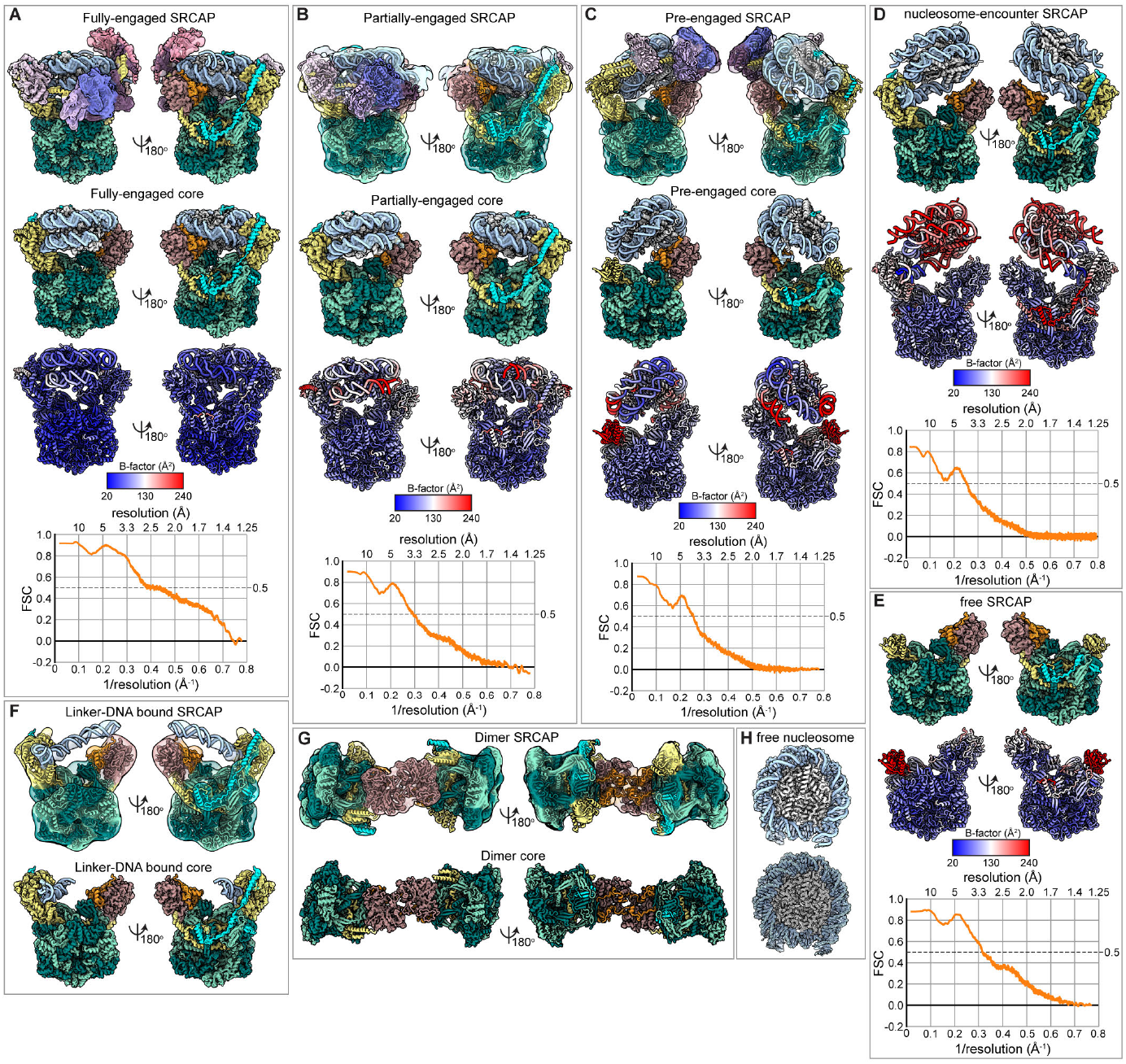
Model building of SRCAP cryo-EM structures. **(A-C)** Overall fit of SRCAP model (A, Fully-engaged; B, Partially-engaged; C, Pre-engaged) into the lower-resolution full cryo-EM map (top). Overall fit of refined SRCAP model (A, Fully-engaged; B, Partially-engaged; C, Pre-engaged) into the high-resolution composite cryo-EM map (middle). Refined model colored by PHENIX B-factor estimation (bottom). Map vs. model Fourier shell correlation (FSC) shown below. **(D-E)** Overall fit of refined SRCAP model (D, nucleosome-encounter; E, free) into the high-resolution composite cryo-EM map (top). Refined model colored by PHENIX B-factor estimation (bottom). Map vs. model Fourier shell correlation (FSC) shown below. **(F-G)** Overall fit of SRCAP model (F, Linker-DNA bound; G, Dimer) into the lower-resolution full cryo-EM map (top). Overall fit of core SRCAP model (F, Linker-DNA bound; G, Dimer) into the high-resolution composite cryo-EM map (bottom). **(H)** Overall fit of free nucleosome model (shown as cartoon, top; and atoms, bottom) into the high resolution cryo-EM map

**Figure S4:**
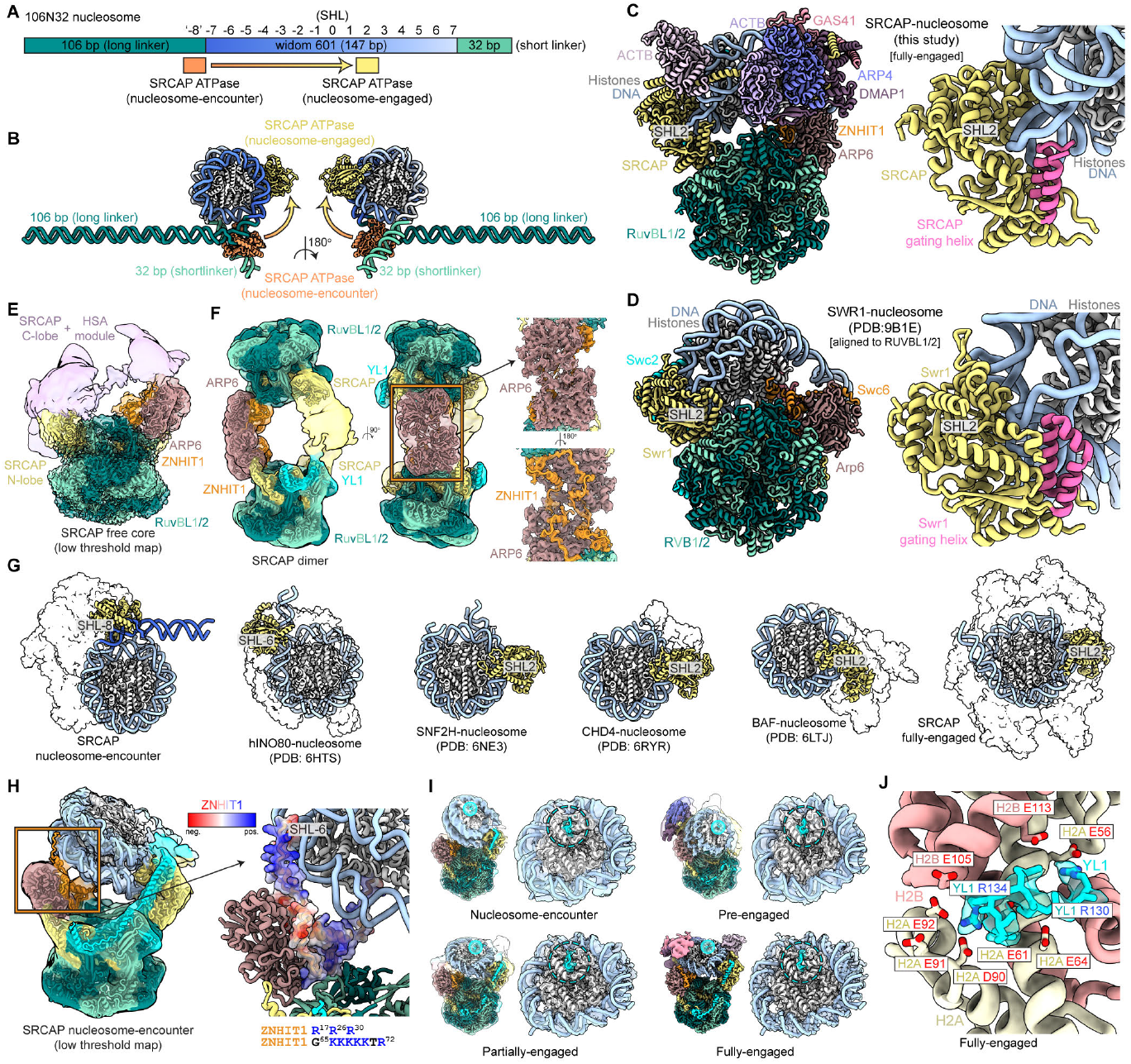
Additional structural analyses of SRCAP. **(A)** Diagram depicting SHL positions of the 106N32 nucleosome used in this study. **(B)** Structural model depicting the location of the SRCAP ATPase in the nucleosome-encounter and -engaged states relative to the long and short linkers. **(C)** Structure of fully-engaged SRCAP-nucleosome complex (this study) and isolated view (right) of SRCAP ATPase bound to SHL2. The gating helix is colored pink. **(D)** Structure of SWR1-nucleosome complex (PDB: 9B1E) aligned to the RUVBL1/2 heterohexamer of SRCAP (panel A) and isolated view (right) of SWR1 ATPase bound to SHL2. The gating helix is colored pink. **(E)** Free SRCAP complex map and model at low threshold. The HSA module interaction with ARP6-ZNHIT1 is observed at lower thresholds. **(F)** Dimeric free SRCAP complex map and model in two different views. The high-resolution focused refinement of the ARP6-ZNHIT1 head-to-head interaction is shown (right). **(G)** Comparison of SRCAP nucleosome-encounter (this study) with SRCAP fully-engaged (this study) and published chromatin remodeler structures (hINO80-nucleosome, PDB:6HTS; SNF2H-nucleosome, PDB: 6NE3; CHD4-nucleosome, PDB: 6RYR; BAF-nucleosome, PDB: 6LTJ) **(H)** Cryo-EM map of SRCAP nucleosome-encounter at low threshold showing two positively charged regions (R^17^, R^26^, R^30^ and K^66^KKKKTR^72^) of ZNHIT1 which interact with the nucleosome near SHL-6. ZNHIT1 is shown as a surface representation colored according to Coulombic electrostatic potential (red: -10 kcal/(mol·e), blue: +10 kcal/(mol·e)). **(I)** Cryo-EM maps of SRCAP nucleosome-encounter, pre-engaged, partially-engaged, and fully-engaged states. YL1 interaction with the H2A-H2B acidic patch is highlighted. **(J)** Close-up of YL1 arginine anchors (R^130^ and R^134^) that interact with the H2A-H2B acidic patch in the fully-engaged state. Residues 130-135 of YL1 and the corresponding cryo-EM density is shown. Atoms are colored by elements (red: oxygen, blue: nitrogen).

**Figure S5:**
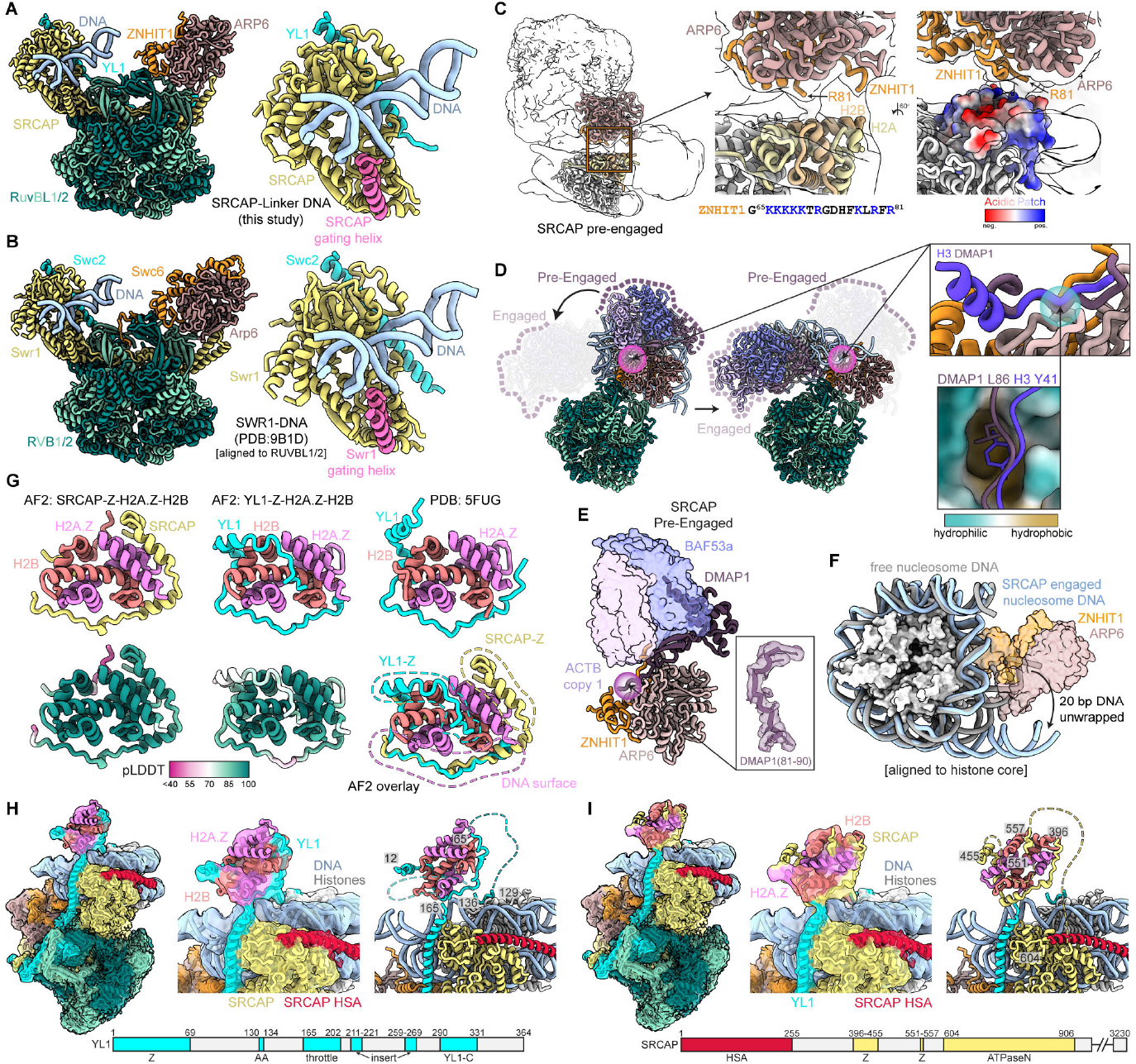
Additional structural analyses of SRCAP. **(A)** Structure of SRCAP bound to linker-DNA (this study) and isolated view of SRCAP ATPase bound to DNA. The gating helix is colored pink. **(B)** Structure of SWR1 bound to DNA (PDB: 9B1D) aligned to the RUVBL1/2 heterohexamer of SRCAP (panel A) and isolated view of SWR1 ATPase bound to DNA. The gating helix is colored pink. **(C)** Cryo-EM map of SRCAP pre-engaged shown as transparent surface showing ZNHIT1 interaction with the H2A-H2B dimer near the acidic patch. The H2A-H2B acidic patch is shown as a surface representation colored according to Coulombic electrostatic potential (red: -10 kcal/(mol·e), blue: +10 kcal/(mol·e)). **(D)** The conformational shift of the HSA module in the transition from pre-engaged to fully-engaged is shown. The ARP6 surface used for binding DMAP1 (pre-engaged) and H3 tail (fully-engaged) are highlighted with a pink sphere. A close-up of the interface is shown in the box with the hydrophobic pocket highlighted with a cyan sphere. The pocket is colored based on the molecular lipophilicity potential (MLP) map with cyan representing hydrophilic and brown representing hydrophobic. **(E)** Isolated view of the A module is shown with the DMAP1 N-terminal region bound by ZNHIT1/ARP6 highlighted with a purple sphere. The cryo-EM density corresponding to DMAP1 residues 81-90 that are bound in the pocket is shown. **(F)** Isolated view of nucleosome, ZNHIT1, ARP6 with free nucleosome DNA as gray and SRCAP engaged nucleosome DNA as light blue. ZNHIT1 and ARP6 are represented as transparent surfaces. The bound DNA (light blue) is unwrapped approximately 20 bp near SHL6 and SHL7. **(G)** Comparison of the crystal structure (PDB: 5FUG) and AlphaFold2-multimer predictions of H2A.Z-H2B chaperone domains of SRCAP complex. The predicted structures are colored by pLDDT confidence score (bottom). The overlay of the YL1 and SRCAP chaperoned H2A.Z-H2B shows an overlapping region chaperoning the DNA interaction surface and unique regions chaperoning the histone interaction surface. **(H)** Putative docking of YL1-H2A.Z-H2B crystal structure (PDB: 5FUG) into low resolution cryo-EM density adjacent to the SRCAP ATPase and HSA helix. Domain map of YL1 is shown below. The chaperone domain is labeled ‘Z’ and arginine anchors as ‘AA’. **(I)** Putative docking of SRCAP-Z-H2A.Z-H2B AlphaFold2-multimer model into low resolution cryo-EM density adjacent to the SRCAP ATPase and HSA helix. Domain map of SRCAP is shown below. The chaperone domains are labeled ‘Z’ and HSA helix as ‘HSA’.

**Figure S6:**
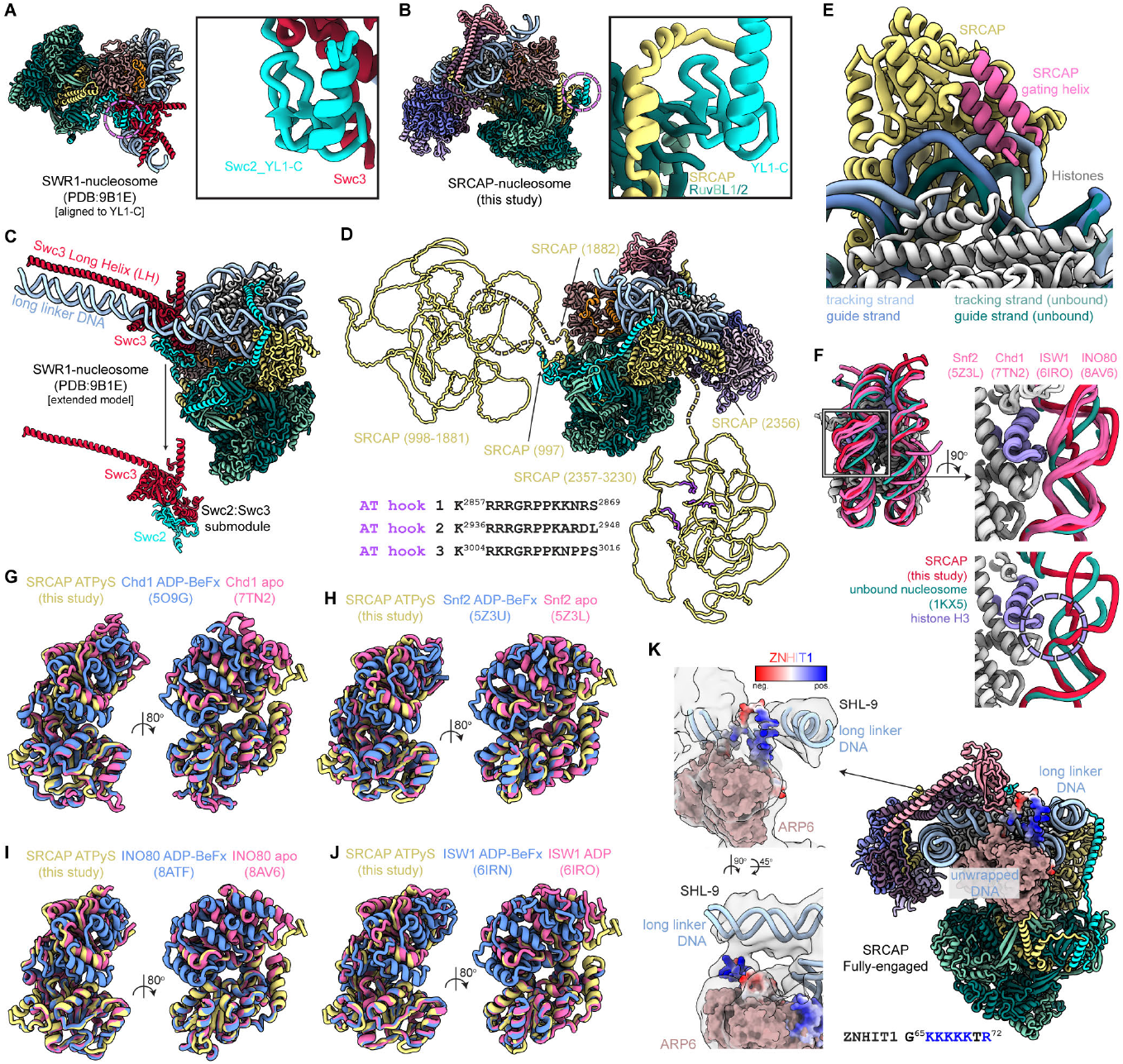
Additional structural analyses of SRCAP. **(A)** SWR1-nucleosome structure (PDB:9B1E) and zoom-in of the Swc2:Swc3 submodule. **(B)** SRCAP-nucleosome structure (this study) and zoom-in of the YL1:SRCAP submodule. **(C)** Extended model of SWR1-nucleosome (PDB:9B1E) highlighting the structured DNA binding elements in yeast Swc2:Swc3 submodule. Swc3 is shown as a surface representation colored according to Coulombic electrostatic potential (red: -10 kcal/(mol·e), blue: +10 kcal/(mol·e)). The yeast Swc2:Swc3 submodule is also shown in isolation (bottom). **(D)** Human YL1:SRCAP submodule lacks structured elements but contains a ∼900 residue intrinsically disordered region (IDR). The C-terminus of SRCAP contains another ∼900 residue IDR that contains three AT hooks (colored purple). **(E)** Binding of SRCAP ATPase induces a large bulge of both tracking and guide strands of DNA. The tracking and guide strands of SRCAP bound DNA are colored light and dark blue, respectively. The tracking and guide strands of free nucleosome DNA are colored light and dark green, respectively. **(F)** Overlay of chromatin remodeler structures (Snf2, PDB: 5Z3L; Chd1, PDB: 7TN2, ISW1, PDB: 6IRO; INO80, PDB: 8AV6; all colored pink), free nucleosome (PDB: 1KX5; colored green), and SRCAP complex (this study; colored red) showing DNA distortion near SHL2.5 where the ATPase binds. SRCAP binding uniquely distorts the DNA and detaches DNA from histone H3 (colored purple). **(G)** Overlay of SRCAP (ATPγS, yellow) with Chd1 closed (ADP-BeFx, blue), and Chd1 open (apo, pink). **(H)** Overlay of SRCAP (ATPγS, yellow) with Snf2 closed (ADP-BeFx, blue), and Snf2 open (apo, pink). **(I)** Overlay of SRCAP (ATPγS, yellow) with INO80 closed (ADP-BeFx, blue), and INO80 open (apo, pink). **(J)** Overlay of SRCAP (ATPγS, yellow) with ISW1 closed (ADP-BeFx, blue), and ISW1 open (ADP, pink). **(K)** The putative location of the highly positive loop of ZNHIT1 in the fully-engaged state is shown. The cryo-EM map is shown transparent and ZHHIT1 is colored according to Coulombic electrostatic potential (red: -10 kcal/(mol·e), blue: +10 kcal/(mol·e)).

**Figure S7:**
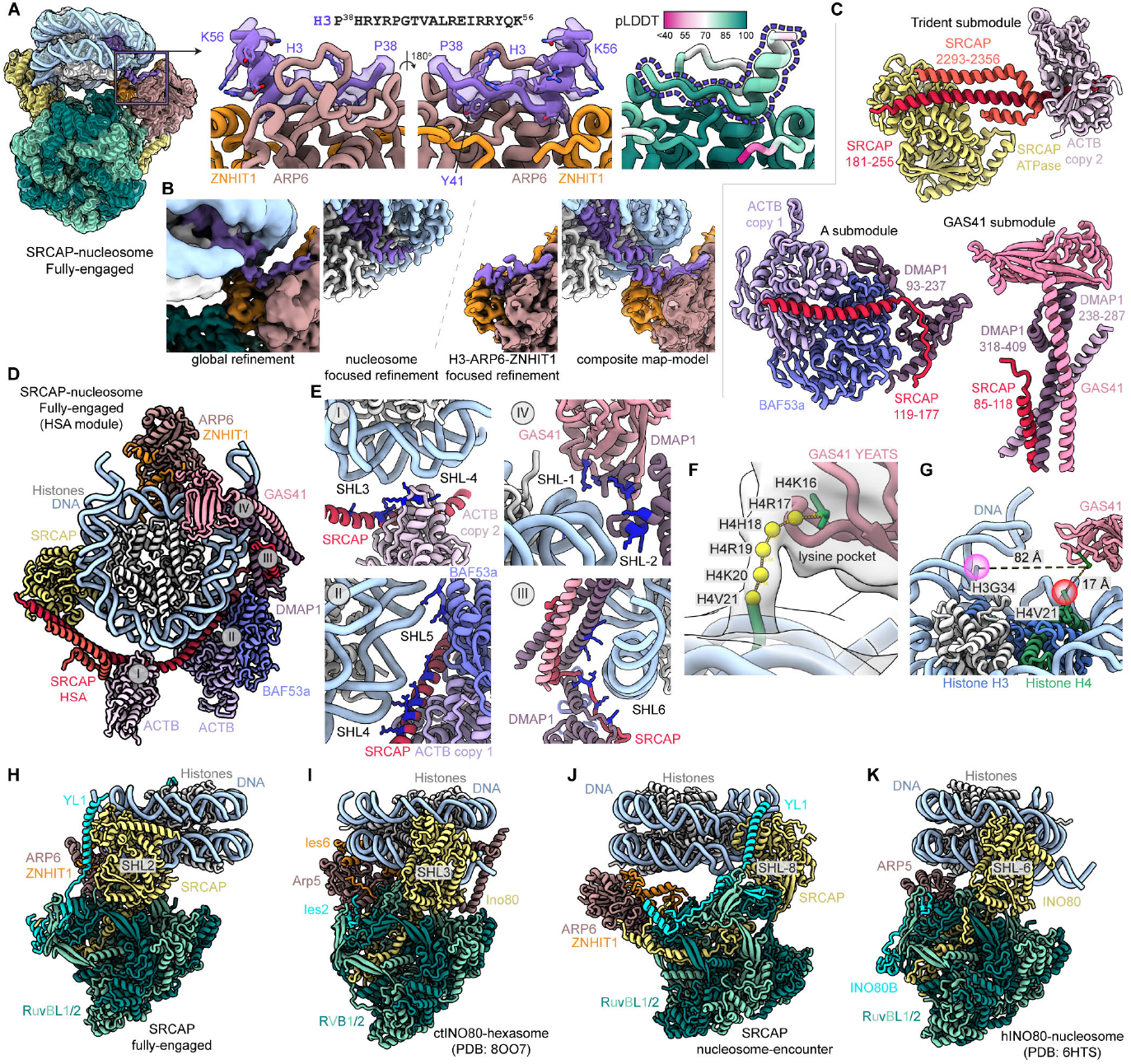
Additional structural analyses of SRCAP. **(A)** Structural details of histone H3 bound by ARP6-ZNHIT1 subunits of SRCAP. The cryo-EM map and model of bound H3 is shown. The AlphaFold2-multimer prediction used as a starting model is shown and colored by pLDDT confidence score. **(B)** The global refinement map, nucleosome focused refinement map, and H3-ARP6-ZNHIT1 focused refinement map show that the H3-ARP6 interaction is dynamic and unable to be fully separated within the fully-engaged state particles. The composite map-model (bottom right) show the two potential positions of the N-terminal alpha-helix of H3 (bound and unbound). **(C)** Isolated views of the three submodules of the HSA module; Trident submodule (top), A module (bottom left), and Gas41 module (bottom right). **(D)** Cryo-EM structure of SRCAP showing HSA module envelopment of the nucleosome. The regions zoomed in are labeled with roman numerals. **(E)** Zoom-in views of SRCAP complex subunits contacting nucleosome DNA. Lysine and arginine residues are colored blue. **(F)** Cryo-EM map of the H4 tail binding to GAS41 YEATS domain. The distance indicates H4K16 is bound in the lysine pocket of YEATS domain. **(G)** Structure of SRCAP fully-engaged showing the YEATS domain and the linear distance from H3 tail (H3R^40^, pink) and H4 tail (H4V^21^, red). **(H)** SRCAP nucleosome fully-engaged structure (this study). The primary ATPase contact site (SHL2) is labeled. **(I)** ctINO80 hexasome structure (PDB: 8OO7). The structure was aligned with the histone core relative to SRCAP nucleosome fully-engaged (panel H). The primary ATPase contact site (SHL3) is labeled. **(J)** SRCAP nucleosome-encounter structure (this study). The primary ATPase contact site (SHL-8) is labeled. **(K)** hINO80 hexasome structure (PDB: 6HTS). The structure was aligned with the histone core relative to SRCAP nucleosome-encounter (panel J). The primary ATPase contact site (SHL-6) is labeled.

**Figure S8:**
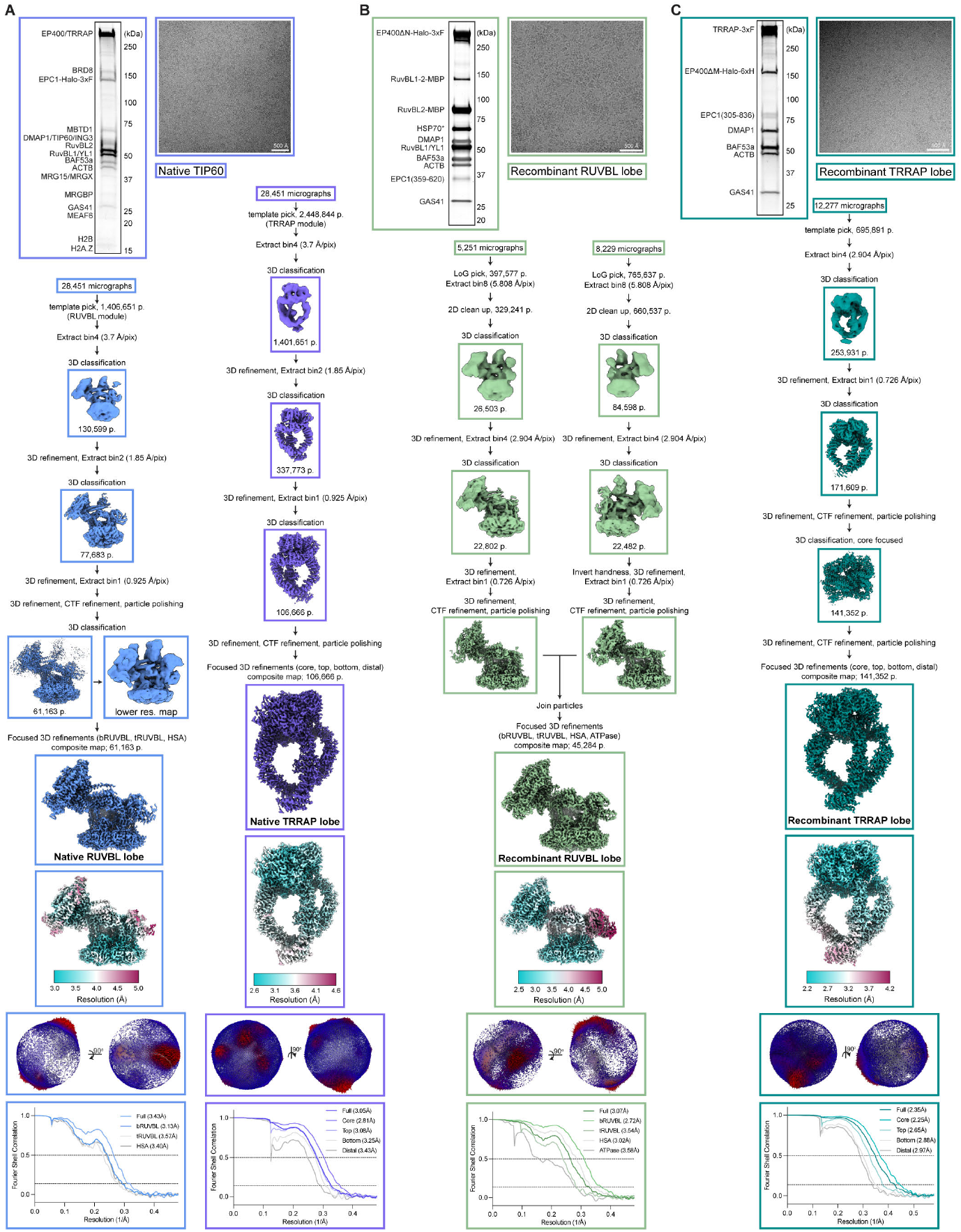
Cryo-EM processing of SRCAP (native TIP60, RUVBL lobe, TRRAP lobe). **(A-C)** SDS-PAGE gel, cryo-EM micrograph, and cryo-EM data processing of native TIP60 (a), ectopically expressed RUVBL lobe (b), and ectopically expressed TRRAP lobe (c). The post processed composite map, local resolution colored map, mask corrected gold-standard Fourier shell correlation (FSC) plots, and orientation distribution plots for global refinements are shown for each state.

**Figure S9:**
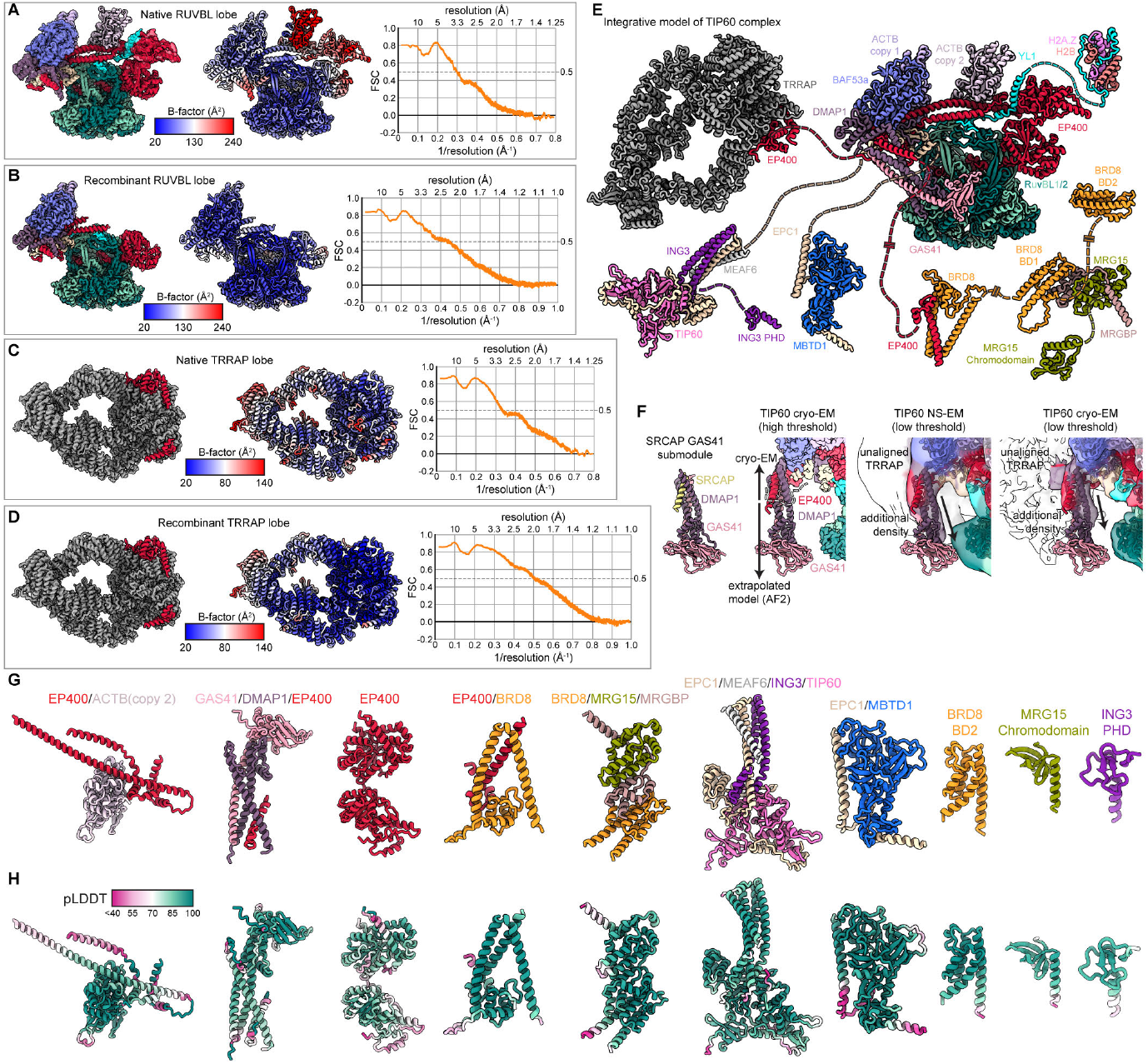
Model building of TIP60 cryo-EM structures and AlphaFold2 predictions. **(A-D)** Overall fit of TIP60 model (a, Native RUVBL lobe; b, Recombinant RUVBL lobe; c, Native TRRAP lobe; d, Recombinant TRRAP lobe) into the high-resolution composite cryo-EM map (left). Refined model colored by PHENIX B-factor estimation (middle). Map vs. model Fourier shell correlation (FSC) is shown (right). **(E)** Full integrative structure of TIP60 generated with cryo-EM structures and AlphaFold2 multimer predictions. **(F)** Extrapolated extension of the EP400-DMAP1-GAS41 submodule in the TIP60 complex. The SRCAP-DMAP1-GAS41 submodule is shown as comparison. The tetrameric coiled-coil structure was extended based on lower resolution negative stain and cryo-EM map. **(G)** AlphaFold2-multimer predictions of various modules of the TIP60 complex. **(H)** Predicted structures in panel A colored by pLDDT confidence score.

**Figure S10:**
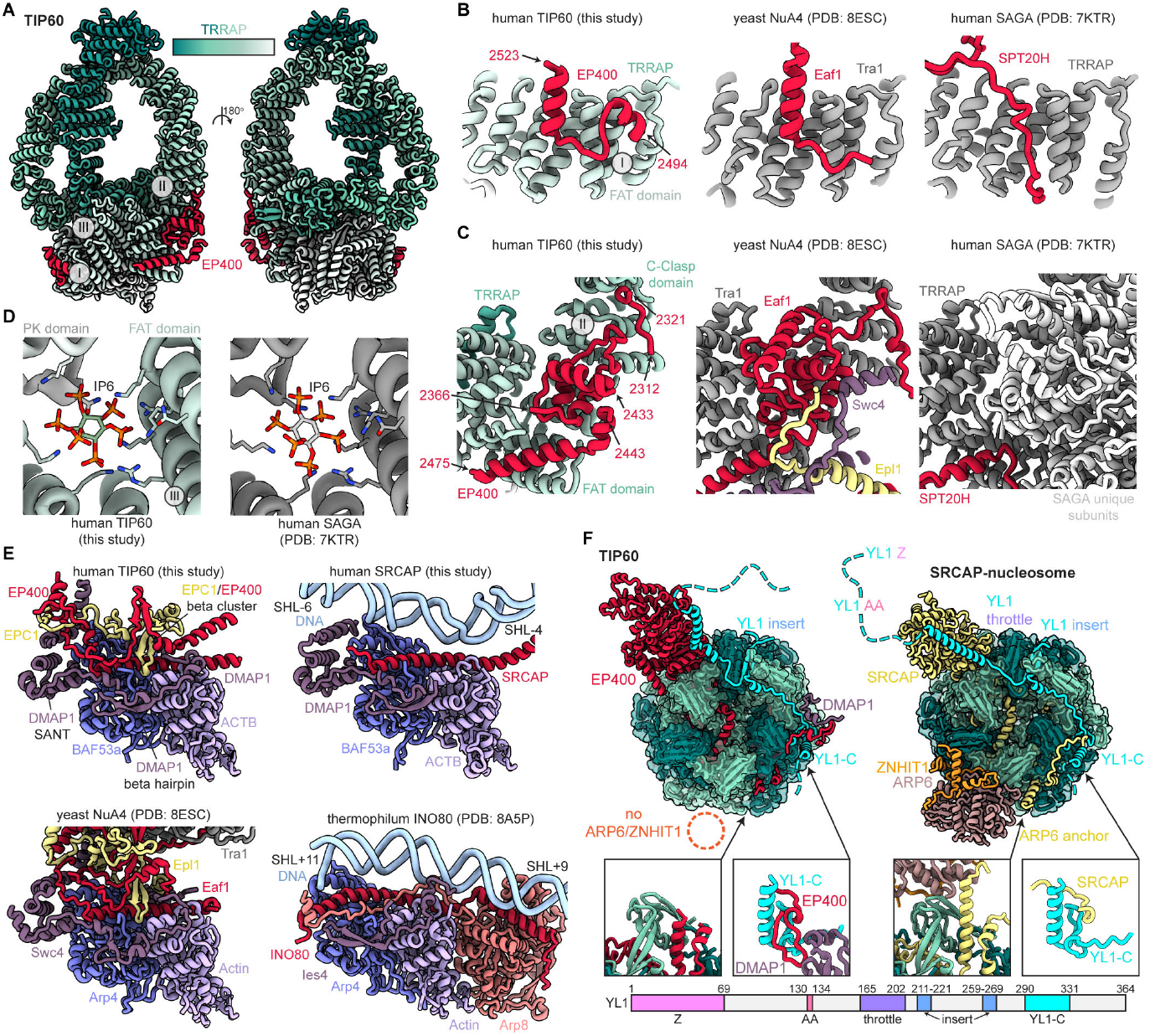
Additional structural analyses of TIP60. **(A)** Cryo-EM structure of TIP60 TRRAP lobe. EP400 is colored red and TRRAP is colored green to white, sequentially. The roman numerals indicate regions focused in subsequent panels. **(B)** Zoom in view of region I of human TIP60 (this study), yeast NuA4 (PDB: 8ESC), and human SAGA (PDB: 7KTR). The structures were aligned to the TRRAP core. EP400 residues 2494-2523 interact with the FAT domain of TRRAP. Tra1/TRRAP interacting subunits (EP400, Eaf1, SPT20H) are colored red. **(C)** Zoom in view of region II of human TIP60, yeast NuA4, and human SAGA. The structures were aligned to the TRRAP core. EP400 residues 2312-2321, 2366-2433, and 2443-2475 interact with the FAT and C-Clasp domain of TRRAP. Tra1/TRRAP interacting subunits (EP400, Eaf1, SPT20H) are colored red. **(D)** Zoom in view of region III of human TIP60 and human SAGA. The structures were aligned to the TRRAP core. Both TIP60 and SAGA contain inositol hexakisphosphate (IP6) between the FAT and PK domain of TRRAP. **(E)** Comparison of the A module of human TIP60 (this study), yeast NuA4 (PDB: 8ESC), human SRCAP (this study), and thermophilum INO80 (PDB: 8A5P). Subunits colored based on homology. Only relevant subunits are shown. **(F)** Top down isolated view of TIP60 RUVBL lobe and SRCAP-nucleosome complex. Only relevant subunits are shown for clarity. Boxes show zoomed in side views of indicated regions. Dotted circle represents absence of ARP6/ZNHIT1 (orange). Dotted lines represent linker between domains of YL1 (cyan) with YL1 Z being the H2A.Z-H2B chaperone domain and YL1 AA being the arginine anchors. A domain map of YL1 is shown at the bottom.

## References

1. Lai, W.K.M., and Pugh, B.F. (2017). Understanding nucleosome dynamics and their links to gene expression and DNA replication. Nat. Rev. Mol. Cell Biol. 18, 548–562.

2. Hauer, M.H., and Gasser, S.M. (2017). Chromatin and nucleosome dynamics in DNA damage and repair. Genes Dev. 31, 2204–2221.

3. Arents, G., Burlingame, R.W., Wang, B.C., Love, W.E., and Moudrianakis, E.N. (1991). The nucleosomal core histone octamer at 3.1 A resolution: a tripartite protein assembly and a left-handed superhelix. Proc. Natl. Acad. Sci. U. S. A. 88, 10148–10152.

4. Luger, K., Mäder, A.W., Richmond, R.K., Sargent, D.F., and Richmond, T.J. (1997). Crystal structure of the nucleosome core particle at 2.8 Å resolution. Nature 389, 251–260.

5. Martire, S., and Banaszynski, L.A. (2020). The roles of histone variants in fine-tuning chromatin organization and function. Nat. Rev. Mol. Cell Biol. 21, 522–541.

6. Ghiraldini, F.G., Filipescu, D., and Bernstein, E. (2021). Solid tumours hijack the histone variant network. Nat. Rev. Cancer 21, 257–275.

7. Giaimo, B.D., Ferrante, F., Herchenröther, A., Hake, S.B., and Borggrefe, T. (2019). The histone variant H2A.Z in gene regulation. Epigenetics Chromatin 12, 37.

8. Colino-Sanguino, Y., Clark, S.J., and Valdes-Mora, F. (2022). The H2A.Z-nucleosome code in mammals: emerging functions. Trends Genet. 38, 273–289.

9. Ruhl, D.D., Jin, J., Cai, Y., Swanson, S., Florens, L., Washburn, M.P., Conaway, R.C., Conaway, J.W., and Chrivia, J.C. (2006). Purification of a human SRCAP complex that remodels chromatin by incorporating the histone variant H2A.Z into nucleosomes. Biochemistry 45, 5671–5677.

10. Wong, M.M., Cox, L.K., and Chrivia, J.C. (2007). The Chromatin Remodeling Protein, SRCAP, Is Critical for Deposition of the Histone Variant H2A.Z at Promoters*. J. Biol. Chem. 282, 26132–26139.

11. Wichmann, J., Pitt, C., Eccles, S., Garnham, A.L., Li-Wai-Suen, C.S.N., May, R., Allan, E., Wilcox, S., Herold, M.J., Smyth, G.K., et al. (2022). Loss of TIP60 (KAT5) abolishes H2AZ lysine 7 acetylation and causes p53, INK4A, and ARF-independent cell cycle arrest. Cell Death Dis. 13, 627.

12. Janas, J.A., Zhang, L., Luu, J.H., Demeter, J., Meng, L., Marro, S.G., Mall, M., Mooney, N.A., Schaukowitch, K., Ng, Y.H., et al. (2022). Tip60-mediated H2A.Z acetylation promotes neuronal fate specification and bivalent gene activation. Mol. Cell 82, 4627–4646.e14.

13. Mizuguchi, G., Shen, X., Landry, J., Wu, W.-H., Sen, S., and Wu, C. (2004). ATP-driven exchange of histone H2AZ variant catalyzed by SWR1 chromatin remodeling complex. Science 303, 343–348.

14. Yang, Y., Zhang, L., Xiong, C., Chen, J., Wang, L., Wen, Z., Yu, J., Chen, P., Xu, Y., Jin, J., et al. (2022). HIRA complex presets transcriptional potential through coordinating depositions of the histone variants H3.3 and H2A.Z on the poised genes in mESCs. Nucleic Acids Res. 50, 191–206.

15. Ravens, S., Yu, C., Ye, T., Stierle, M., and Tora, L. (2015). Tip60 complex binds to active Pol II promoters and a subset of enhancers and coregulates the c-Myc network in mouse embryonic stem cells. Epigenetics Chromatin 8, 45.

16. Pradhan, S.K., Su, T., Yen, L., Jacquet, K., Huang, C., Côté, J., Kurdistani, S.K., and Carey, M.F. (2016). EP400 Deposits H3.3 into Promoters and Enhancers during Gene Activation. Mol. Cell 61, 27–38.

17. Schones, D.E., Cui, K., Cuddapah, S., Roh, T.-Y., Barski, A., Wang, Z., Wei, G., and Zhao, K. (2008). Dynamic regulation of nucleosome positioning in the human genome. Cell 132, 887–898.

18. Jin, C., Zang, C., Wei, G., Cui, K., Peng, W., Zhao, K., and Felsenfeld, G. (2009). H3.3/H2A.Z double variant-containing nucleosomes mark “nucleosome-free regions” of active promoters and other regulatory regions. Nat. Genet. 41, 941–945.

19. Cole, L., Kurscheid, S., Nekrasov, M., Domaschenz, R., Vera, D.L., Dennis, J.H., and Tremethick, D.J. (2021). Multiple roles of H2A.Z in regulating promoter chromatin architecture in human cells. Nat. Commun. 12, 2524.

20. Cai, Y., Jin, J., Florens, L., Swanson, S.K., Kusch, T., Li, B., Workman, J.L., Washburn, M.P., Conaway, R.C., and Conaway, J.W. (2005). The mammalian YL1 protein is a shared subunit of the TRRAP/TIP60 histone acetyltransferase and SRCAP complexes. J. Biol. Chem. 280, 13665–13670.

21. Jacquet, K., Fradet-Turcotte, A., Avvakumov, N., Lambert, J.-P., Roques, C., Pandita, R.K., Paquet, E., Herst, P., Gingras, A.-C., Pandita, T.K., et al. (2016). The TIP60 Complex Regulates Bivalent Chromatin Recognition by 53BP1 through Direct H4K20me Binding and H2AK15 Acetylation. Mol. Cell 62, 409–421.

22. Willhoft, O., Ghoneim, M., Lin, C.-L., Chua, E.Y.D., Wilkinson, M., Chaban, Y., Ayala, R., McCormack, E.A., Ocloo, L., Rueda, D.S., et al. (2018). Structure and dynamics of the yeast SWR1-nucleosome complex. Science 362. 10.1126/science.aat7716.

23. Louder, R.K. (2024). Molecular basis of global promoter sensing and nucleosome capture by the SWR1 chromatin remodeler.

24. He, S., Wu, Z., Tian, Y., Yu, Z., Yu, J., Wang, X., Li, J., Liu, B., and Xu, Y. (2020). Structure of nucleosome-bound human BAF complex. Science 367, 875–881.

25. Yuan, J., Chen, K., Zhang, W., and Chen, Z. (2022). Structure of human chromatin-remodelling PBAF complex bound to a nucleosome. Nature 605, 166–171.

26. Armache, J.P., Gamarra, N., Johnson, S.L., Leonard, J.D., Wu, S., Narlikar, G.J., and Cheng, Y. (2019). Cryo-EM structures of remodelernucleosome intermediates suggest allosteric control through the nucleosome. Elife 8. 10.7554/eLife.46057.

27. Ayala, R., Willhoft, O., Aramayo, R.J., Wilkinson, M., McCormack, E.A., Ocloo, L., Wigley, D.B., and Zhang, X. (2018). Structure and regulation of the human INO80-nucleosome complex. Nature 556, 391–395.

28. Farnung, L., Ochmann, M., and Cramer, P. (2020). Nucleosome-CHD4 chromatin remodeler structure maps human disease mutations. Elife 9. 10.7554/eLife.56178.

29. Eustermann, S., Patel, A.B., Hopfner, K.-P., He, Y., and Korber, P. (2023). Energy-driven genome regulation by ATP-dependent chromatin remodellers. Nat. Rev. Mol. Cell Biol. 10.1038/s41580-023-00683-y.

30. Zukin, S.A., Marunde, M.R., Popova, I.K., Soczek, K.M., Nogales, E., and Patel, A.B. (2022). Structure and flexibility of the yeast NuA4 histone acetyltransferase complex. Elife 11. 10.7554/eLife.81400.

31. Carcamo, C.C., Poyton, M.F., Ranjan, A., Park, G., Louder, R.K., Feng, X.A., Kim, J.M., Dzu, T., Wu, C., and Ha, T. (2022). ATP binding facilitates target search of SWR1 chromatin remodeler by promoting one-dimensional diffusion on DNA. Elife 11. 10.7554/eLife.77352.

32. Liang, X., Shan, S., Pan, L., Zhao, J., Ranjan, A., Wang, F., Zhang, Z., Huang, Y., Feng, H., Wei, D., et al. (2016). Structural basis of H2A.Z recognition by SRCAP chromatin-remodeling subunit YL1. Nat. Struct. Mol. Biol. 23, 317–323.

33. Latrick, C.M., Marek, M., Ouararhni, K., Papin, C., Stoll, I., Ignatyeva, M., Obri, A., Ennifar, E., Dimitrov, S., Romier, C., et al. (2016). Molecular basis and specificity of H2A.Z-H2B recognition and deposition by the histone chaperone YL1. Nat. Struct. Mol. Biol. 23, 309–316.

34. Hong, J., Feng, H., Wang, F., Ranjan, A., Chen, J., Jiang, J., Ghirlando, R., Xiao, T.S., Wu, C., and Bai, Y. (2014). The catalytic subunit of the SWR1 remodeler is a histone chaperone for the H2A.Z-H2B dimer. Mol. Cell 53, 498–505.

35. Poyton, M.F., Feng, X.A., Ranjan, A., Lei, Q., Wang, F., Zarb, J.S., Louder, R.K., Park, G., Jo, M.H., Ye, J., et al. (2022). Coordinated DNA and histone dynamics drive accurate histone H2A.Z exchange. Sci Adv 8, eabj5509.

36. Patil, A., Strom, A.R., Paulo, J.A., Collings, C.K., Ruff, K.M., Shinn, M.K., Sankar, A., Cervantes, K.S., Wauer, T., St Laurent, J.D., et al. (2023). A disordered region controls cBAF activity via condensation and partner recruitment. Cell 186, 4936–4955.e26.

37. Greenberg, R.S., Long, H.K., Swigut, T., and Wysocka, J. (2019). Single Amino Acid Change Underlies Distinct Roles of H2A.Z Subtypes in Human Syndrome. Cell 178, 1421–1436.e24.

38. Yan, L., Wu, H., Li, X., Gao, N., and Chen, Z. (2019). Structures of the ISWI-nucleosome complex reveal a conserved mechanism of chromatin remodeling. Nat. Struct. Mol. Biol. 26, 258–266.

39. Li, M., Xia, X., Tian, Y., Jia, Q., Liu, X., Lu, Y., Li, M., Li, X., and Chen, Z. (2019). Mechanism of DNA translocation underlying chromatin remodelling by Snf2. Nature 567, 409–413.

40. Farnung, L., Vos, S.M., Wigge, C., and Cramer, P. (2017). Nucleosome– Chd1 structure and implications for chromatin remodelling. Nature 550, 539–542.

41. Nodelman, I.M., Das, S., Faustino, A.M., Fried, S.D., Bowman, G.D., and Armache, J.-P. (2022). Nucleosome recognition and DNA distortion by the Chd1 remodeler in a nucleotide-free state. Nat. Struct. Mol. Biol. 29, 121–129.

42. Kunert, F., Metzner, F.J., Jung, J., Höpfler, M., Woike, S., Schall, K., Kostrewa, D., Moldt, M., Chen, J.-X., Bantele, S., et al. (2022). Structural mechanism of extranucleosomal DNA readout by the INO80 complex. Sci Adv 8, eadd3189.

43. Armeev, G.A., Kniazeva, A.S., Komarova, G.A., Kirpichnikov, M.P., and Shaytan, A.K. (2021). Histone dynamics mediate DNA unwrapping and sliding in nucleosomes. Nat. Commun. 12, 1–15.

44. Hsu, C.-C., Shi, J., Yuan, C., Zhao, D., Jiang, S., Lyu, J., Wang, X., Li, H., Wen, H., Li, W., et al. (2018). Recognition of histone acetylation by the GAS41 YEATS domain promotes H2A.Z deposition in non-small cell lung cancer. Genes Dev. 32, 58–69.

45. Cho, H.J., Li, H., Linhares, B.M., Kim, E., Ndoj, J., Miao, H., Grembecka, J., and Cierpicki, T. (2018). GAS41 Recognizes Diacetylated Histone H3 through a Bivalent Binding Mode. ACS Chem. Biol. 13, 2739–2746.

46. Kikuchi, M., Morita, S., Goto, M., Wakamori, M., Katsura, K., Hanada, K., Shirouzu, M., and Umehara, T. (2022). Elucidation of binding preferences of YEATS domains to site-specific acetylated nucleosome core particles. J. Biol. Chem. 298, 102164.

47. Kikuchi, M., Takase, S., Konuma, T., Noritsugu, K., Sekine, S., Ikegami, T., Ito, A., and Umehara, T. (2023). GAS41 promotes H2A.Z deposition through recognition of the N terminus of histone H3 by the YEATS domain. Proc. Natl. Acad. Sci. U. S. A. 120, e2304103120.

48. Lalonde, M.-E., Cheng, X., and Côté, J. (2014). Histone target selection within chromatin: an exemplary case of teamwork. Genes Dev. 28, 1029–1041.

49. Marunde, M.R., Fuchs, H.A., Burg, J.M., Popova, I.K., Vaidya, A., Hall, N.W., Weinzapfel, E.N., Meiners, M.J., Watson, R., Gillespie, Z.B., et al. (2024). Nucleosome conformation dictates the histone code. Elife 13. 10.7554/eLife.78866.

50. Shvedunova, M., and Akhtar, A. (2022). Modulation of cellular processes by histone and non-histone protein acetylation. Nat. Rev. Mol. Cell Biol. 23, 329–349.

51. Kusch, T., Florens, L., Macdonald, W.H., Swanson, S.K., Glaser, R.L., Yates, J.R., 3rd, Abmayr, S.M., Washburn, M.P., and Workman, J.L. (2004). Acetylation by Tip60 is required for selective histone variant exchange at DNA lesions. Science 306, 2084–2087.

52. Gévry, N., Chan, H.M., Laflamme, L., Livingston, D.M., and Gaudreau, L. (2007). p21 transcription is regulated by differential localization of histone H2A.Z. Genes Dev. 21, 1869–1881.

53. Park, J.H., Sun, X.-J., and Roeder, R.G. (2010). The SANT domain of p400 ATPase represses acetyltransferase activity and coactivator function of TIP60 in basal p21 gene expression. Mol. Cell. Biol. 30, 2750–2761.

54. Zhang, H., Devoucoux, M., Song, X., Li, L., Ayaz, G., Cheng, H., Tempel, W., Dong, C., Loppnau, P., Côté, J., et al. (2020). Structural Basis for EPC1-Mediated Recruitment of MBTD1 into the NuA4/TIP60 Acetyltransferase Complex. Cell Rep. 30, 3996–4002.e4.

55. Herbst, D.A., Esbin, M.N., Louder, R.K., Dugast-Darzacq, C., Dailey, G.M., Fang, Q., Darzacq, X., Tjian, R., and Nogales, E. (2021). Structure of the human SAGA coactivator complex. Nat. Struct. Mol. Biol. 28, 989–996.

56. Mirdita, M., Schütze, K., Moriwaki, Y., Heo, L., Ovchinnikov, S., and Steinegger, M. (2022). ColabFold: making protein folding accessible to all. Nat. Methods 19, 679–682.

57. Evans, R., O’Neill, M., Pritzel, A., Antropova, N., Senior, A., Green, T., Žídek, A., Bates, R., Blackwell, S., Yim, J., et al. (2021). Protein complex prediction with AlphaFold-Multimer. bioRxiv. 10.1101/2021.10.04.463034.

58. Jumper, J., Evans, R., Pritzel, A., Green, T., Figurnov, M., Ronneberger, O., Tunyasuvunakool, K., Bates, R., Žídek, A., Potapenko, A., et al. (2021). Highly accurate protein structure prediction with AlphaFold. Nature 596, 583–589.

59. Xie, T., Zmyslowski, A.M., Zhang, Y., and Radhakrishnan, I. (2015). Structural Basis for Multi-specificity of MRG Domains. Structure 23, 1049–1057.

60. Murr, R., Vaissière, T., Sawan, C., Shukla, V., and Herceg, Z. (2007). Orchestration of chromatin-based processes: mind the TRRAP. Oncogene 26, 5358–5372.

61. von Hippel, P.H., and Berg, O.G. (1989). Facilitated target location in biological systems. J. Biol. Chem. 264, 675–678.

62. Mirny, L., Slutsky, M., Wunderlich, Z., Tafvizi, A., Leith, J., and Kosmrlj, A. (2009). How a protein searches for its site on DNA: the mechanism of facilitated diffusion. J. Phys. A: Math. Theor. 42, 434013.

63. Kim, J.M., Carcamo, C.C., Jazani, S., Xie, Z., Feng, X.A., Poyton, M., Holland, K.L., Grimm, J.B., Lavis, L.D., Ha, T., et al. (2023). Dynamic 1D Search and Processive Nucleosome Translocations by RSC and ISW2 Chromatin Remodelers. bioRxiv, 2023.06.13.544671. 10.1101/2023.06.13.544671.

64. Luk, E., Ranjan, A., Fitzgerald, P.C., Mizuguchi, G., Huang, Y., Wei, D., and Wu, C. (2010). Stepwise histone replacement by SWR1 requires dual activation with histone H2A.Z and canonical nucleosome. Cell 143, 725–736.

65. Ranjan, A., Wang, F., Mizuguchi, G., Wei, D., Huang, Y., and Wu, C. (2015). H2A histone-fold and DNA elements in nucleosome activate SWR1-mediated H2A.Z replacement in budding yeast. Elife 4, e06845.

66. Suto, R.K., Clarkson, M.J., Tremethick, D.J., and Luger, K. (2000). Crystal structure of a nucleosome core particle containing the variant histone H2A.Z. Nat. Struct. Biol. 7, 1121–1124.

67. Eustermann, S., Schall, K., Kostrewa, D., Lakomek, K., Strauss, M., Moldt, M., and Hopfner, K.-P. (2018). Structural basis for ATP-dependent chromatin remodelling by the INO80 complex. Nature 556, 386–390.

68. Zhang, M., Jungblut, A., Kunert, F., Hauptmann, L., Hoffmann, T., Kolesnikova, O., Metzner, F., Moldt, M., Weis, F., DiMaio, F., et al. (2023). Hexasome-INO80 complex reveals structural basis of noncanonical nucleosome remodeling. Science 381, 313–319.

69. Wu, H., Muñoz, E.N., Hsieh, L.J., Chio, U.S., Gourdet, M.A., Narlikar, G.J., and Cheng, Y. (2023). Reorientation of INO80 on hexasomes reveals basis for mechanistic versatility. Science 381, 319–324.

70. McGinty, R.K., and Tan, S. (2021). Principles of nucleosome recognition by chromatin factors and enzymes. Curr. Opin. Struct. Biol. 71, 16–26.

71. Yu, J., Sui, F., Gu, F., Li, W., Yu, Z., Wang, Q., He, S., Wang, L., and Xu, Y. (2024). Structural insights into histone exchange by human SRCAP complex. Cell Discov 10, 15.

72. Scacchetti, A., Schauer, T., Reim, A., Apostolou, Z., Campos Sparr, A., Krause, S., Heun, P., Wierer, M., and Becker, P.B. (2020). Drosophila SWR1 and NuA4 complexes are defined by DOMINO isoforms. Elife 9, e56325.

73. Zhao, B., Chen, Y., Jiang, N., Yang, L., Sun, S., Zhang, Y., Wen, Z., Ray, L., Liu, H., Hou, G., et al. (2019). Znhit1 controls intestinal stem cell maintenance by regulating H2A.Z incorporation. Nat. Commun. 10, 1071.

74. Cuadrado, A., Corrado, N., Perdiguero, E., Lafarga, V., Muñoz-Canoves, P., and Nebreda, A.R. (2010). Essential role of p18Ham-let/SRCAP-mediated histone H2A.Z chromatin incorporation in muscle differentiation. EMBO J. 29, 2014–2025.

75. Sun, S., Jiang, Y., Zhang, Q., Pan, H., Li, X., Yang, L., Huang, M., Wei, W., Wang, X., Qiu, M., et al. (2022). Znhit1 controls meiotic initiation in male germ cells by coordinating with Stra8 to activate meiotic gene expression. Dev. Cell 57, 901–913.e4.

76. Börner, K., and Becker, P.B. (2016). Splice variants of the SWR1-type nucleosome remodeling factor Domino have distinct functions during Drosophila melanogaster oogenesis. Development 143, 3154–3167.

77. Scacchetti, A., and Becker, P.B. (2021). Variation on a theme: Evolutionary strategies for H2A.Z exchange by SWR1-type remodelers. Curr. Opin. Cell Biol. 70, 1–9.

78. Abmayr, S.M., Yao, T., Parmely, T., and Workman, J.L. (2006). Preparation of nuclear and cytoplasmic extracts from mammalian cells. Curr. Protoc. Mol. Biol. Chapter 12, Unit 12.1.

79. Dyer, P.N., Edayathumangalam, R.S., White, C.L., Bao, Y., Chakravarthy, S., Muthurajan, U.M., and Luger, K. (2004). Reconstitution of nucleosome core particles from recombinant histones and DNA. Methods Enzymol. 375, 23–44.

80. Juven-Gershon, T., Cheng, S., and Kadonaga, J.T. (2006). Rational design of a super core promoter that enhances gene expression. Nat. Methods 3, 917–922.

81. Lowary, P.T., and Widom, J. (1998). New DNA sequence rules for high affinity binding to histone octamer and sequence-directed nucleosome positioning. J. Mol. Biol. 276, 19–42.

82. Punjani, A., Rubinstein, J.L., Fleet, D.J., and Brubaker, M.A. (2017). cryoSPARC: algorithms for rapid unsupervised cryo-EM structure determination. Nat. Methods 14, 290–296.

83. Patel, A., Toso, D., Litvak, A., and Nogales, E. (2021). Efficient graphene oxide coating improves cryo-EM sample preparation and data collection from tilted grids. bioRxiv, 2021.03.08.434344. 10.1101/2021.03.08.434344.

84. Mastronarde, D.N. (2005). Automated electron microscope tomography using robust prediction of specimen movements. J. Struct. Biol. 152, 36–51.

85. Kimanius, D., Dong, L., Sharov, G., Nakane, T., and Scheres, S.H.W. (2021). New tools for automated cryo-EM single-particle analysis in RELION-4.0. Biochem. J 478, 4169–4185.

86. Zheng, S.Q., Palovcak, E., Armache, J.-P., Verba, K.A., Cheng, Y., and Agard, D.A. (2017). MotionCor2: anisotropic correction of beam-induced motion for improved cryo-electron microscopy. Nat. Methods 14, 331–332.

87. Zhang, K. (2016). Gctf: Real-time CTF determination and correction. J. Struct. Biol. 193, 1–12.

88. Rohou, A., and Grigorieff, N. (2015). CTFFIND4: Fast and accurate defocus estimation from electron micrographs. J. Struct. Biol. 192, 216–221.

89. Henderson, R., Sali, A., Baker, M.L., Carragher, B., Devkota, B., Downing, K.H., Egelman, E.H., Feng, Z., Frank, J., Grigorieff, N., et al. (2012). Outcome of the first electron microscopy validation task force meeting. Structure 20, 205–214.

90. Zivanov, J., Nakane, T., Forsberg, B.O., Kimanius, D., Hagen, W.J., Lindahl, E., and Scheres, S.H. (2018). New tools for automated highresolution cryo-EM structure determination in RELION-3. Elife 7. 10.7554/eLife.42166.

91. Zivanov, J., Nakane, T., and Scheres, S.H.W. (2019). A Bayesian approach to beam-induced motion correction in cryo-EM single-particle analysis. IUCrJ 6, 5–17.

92. Sanchez-Garcia, R., Gomez-Blanco, J., Cuervo, A., Carazo, J.M., Sorzano, C.O.S., and Vargas, J. (2021). DeepEMhancer: a deep learning solution for cryo-EM volume post-processing. Commun Biol 4, 874.

93. Emsley, P., Lohkamp, B., Scott, W.G., and Cowtan, K. (2010). Features and development of Coot. Acta Crystallogr. D Biol. Crystallogr. 66, 486–501.

94. Croll, T.I. (2018). ISOLDE: a physically realistic environment for model building into low-resolution electron-density maps. Acta Crystallogr D Struct Biol 74, 519–530.

95. Meng, E.C., Goddard, T.D., Pettersen, E.F., Couch, G.S., Pearson, Z.J., Morris, J.H., and Ferrin, T.E. (2023). UCSF ChimeraX: Tools for structure building and analysis. Protein Sci. 32, e4792.

96. Liebschner, D., Afonine, P.V., Baker, M.L., Bunkóczi, G., Chen, V.B., Croll, T.I., Hintze, B., Hung, L.W., Jain, S., McCoy, A.J., et al. (2019). Macromolecular structure determination using X-rays, neutrons and electrons: recent developments in Phenix. Acta Crystallogr D Struct Biol 75, 861–877.

97. Afonine, P.V., Klaholz, B.P., Moriarty, N.W., Poon, B.K., Sobolev, O.V., Terwilliger, T.C., Adams, P.D., and Urzhumtsev, A. (2018). New tools for the analysis and validation of cryo-EM maps and atomic models. Acta Crystallogr D Struct Biol 74, 814–840.

98. Williams, C.J., Headd, J.J., Moriarty, N.W., Prisant, M.G., Videau, L.L., Deis, L.N., Verma, V., Keedy, D.A., Hintze, B.J., Chen, V.B., et al. (2018). MolProbity: More and better reference data for improved allatom structure validation. Protein Sci. 27, 293–315.

99. Goddard, T.D., Huang, C.C., Meng, E.C., Pettersen, E.F., Couch, G.S., Morris, J.H., and Ferrin, T.E. (2018). UCSF ChimeraX: Meeting modern challenges in visualization and analysis. Protein Sci. 27, 14–25.

100. Hammond, C.M., Strømme, C.B., Huang, H., Patel, D.J., and Groth, A. (2017). Histone chaperone networks shaping chromatin function. Nat. Rev. Mol. Cell Biol. 18, 141–158.

101. Racki, L.R., Yang, J.G., Naber, N., Partensky, P.D., Acevedo, A., Purcell, T.J., Cooke, R., Cheng, Y., and Narlikar, G.J. (2009). The chromatin remodeller ACF acts as a dimeric motor to space nucleosomes. Nature 462, 1016–1021.

102. Willhoft, O., McCormack, E.A., Aramayo, R.J., Bythell-Douglas, R., Ocloo, L., Zhang, X., and Wigley, D.B. (2017). Crosstalk within a functional INO80 complex dimer regulates nucleosome sliding. Elife 6. 10.7554/eLife.25782.

103. Sun, L., Pierrakeas, L., Li, T., and Luk, E. (2020). Thermosensitive Nucleosome Editing Reveals the Role of DNA Sequence in Targeted Histone Variant Deposition. Cell Rep. 30, 257–268.e5.

104. Rhee, H.S., Bataille, A.R., Zhang, L., and Pugh, B.F. (2014). Subnucleosomal structures and nucleosome asymmetry across a genome. Cell 159, 1377–1388.

